# Neuronal heparan sulfate controls the brain accumulation and glymphatic clearance of pathogenic α-synuclein

**DOI:** 10.64898/2026.11.25.769298

**Authors:** Saumya Digraskar, Piyali Das, Ryan Bash, Jaewon Huh, Rodrigo Queiroz Guimaraes, Hailey Houson, Emily E Helman, Sharon L Samuel, Geir Sajdak, Charles Murchison, Marla Gearing, Jeffrey Iliff, Jeffrey Esko, Rita Cowell, Suzanne E Lapi, Anna Sorace, Laura Volpicelli-Daley, Patricia Aguilar-Calvo

## Abstract

In Parkinson’s disease and dementia with Lewy bodies, fibrillar aggregates of the α-synuclein (α-syn) protein are released into the extracellular space and drive disease progression, yet the mechanisms governing their in vivo neuronal uptake and clearance remain poorly understood. Heparan sulfate proteoglycans (HSPGs) act as receptors for α-syn fibril internalization in iPSC-derived dopaminergic neurons and could therefore advance synucleinopathy progression in patients. Here we show that α-syn fibrils interact electrostatically with the sulfate groups in the HS chains with high affinity, and that genetically reducing the sulfation of neuronal HS reduces the accumulation of α-syn aggregates in multiple brain areas, the loss of dopaminergic neurons, and the development of a locomotor behavioral phenotype in mice intracerebrally injected with α-syn recombinant fibrils (PFFs). In vivo imaging revealed that HS is also a molecular obstacle to the glymphatic clearance of α-syn fibrils, as radiolabeled α-syn PFFs travel faster through the brain towards cerebrospinal clearance in mice with less sulfated neuronal HS than in wild-type mice. Collectively, this study provides in vivo evidence that neuronal HS accelerates α-syn pathology, potentially by promoting neuronal uptake while slowing the glymphatic clearance of α-syn fibrils and, therefore, is a promising therapeutic target to halt synucleinopathies.

## Introduction

Parkinson’s disease (PD) and dementia with Lewy bodies (DLB) are progressive neurodegenerative disorders defined by the intracellular accumulation of α-synuclein (α-syn) aggregates^1^. In these diseases, Lewy bodies (LB) and Lewy neurites, comprised mostly of aggregated α-syn in neurons, disrupt cell’s homeostasis and lead to neuronal dysfunction and loss^2,3^. α-Syn pathology shows a predictable spatio-temporal progression in PD and DLB patients and in mouse models of synucleinopathy, suggesting that α-syn aggregates may propagate along axons and synapses^4–7^. However, pathological α-syn is also detected in extracellular fluids of PD and DLB patients, such as cerebrospinal fluid (CSF) and blood^8–12^, and α-syn inclusions are not always confined to anatomically connected neurons^13^. Intracerebral injections of anti-α-syn antibodies slow pathology in PD mouse models^14^, and blocking the clearance of extracellular α-syn fibrils either into the CSF via the glymphatic system or from the CSF to the periphery via the meningeal lymphatic drainage accelerates the disease^15,16^. Moreover, recent studies proposed the extracellular transport of α-syn monomers as a fast-track pathway for the propagation of α-syn pathology to remote, anatomically unconnected brain regions^17^. Collectively, mounting evidence suggests that extracellular α-syn aggregates significantly contribute to disease progression and support the design of strategies to impair their neuronal internalization and enhance their glymphatic clearance to slow or halt synucleinopathy progression. Yet only a few studies investigate the extracellular events of synucleinopathies.

Heparan sulfate proteoglycans (HSPGs) are glycoproteins abundant at the neuronal surface, presynaptic terminals, the brain extracellular matrix and the vascular basement membrane^18,19^, where they bind tau, amyloid-β (Aβ), and prion fibrils, promoting their cellular internalization and in vitro fibril formation^20–23^. HSPGs are comprised of a protein core with at least one polysaccharide HS chain attached. The interaction of HSPGs with ligands is determined by the HS chain length and the level and pattern of sulfation^18^. HS is abundant in the plaques of Alzheimer’s disease and prion disease patients^22,24,25^, and genetically shortening or desulfating HS chains reduces the deposition of tau, Aβ, and prions in mouse models of tauopathy and Alzheimer’s disease, and in prion-infected mice^22,23,26,27^. Moreover, reducing HS sulfation accelerates the glymphatic clearance of radio-labeled prion monomers in mice, suggesting that the electrochemical interactions of prions with the HS’ sulfate groups in the brain and spine extracellular space might significantly contribute to prion pathology^23^.

HSPGs mediate the uptake of α-syn fibrils in immortalized cells^19–21,28–30^, mouse hippocampal neurons^31^, and more recently in human iPSC-derived dopaminergic neurons^32^. Moreover, the upregulated expression of GPC6 (the gene encoding the cell-surface glycosylphosphatidylinositol-anchored HSPG glypican 6) in iPSC-derived dopaminergic neurons from individuals with sporadic PD and genetic PD suggests a potential pathologic role of HSPG in synucleinopathies^33^. In this study, we investigated the contribution of neuronal HSPG to α-syn pathology using mice with less sulfated HS in neurons due to conditional depletion of the gene that encodes the N-deacetylase/N-sulfotransferase enzyme, *Ndst1*. We found that mice with less sulfated neuronal HS and intrastriatally injected with α-syn preformed fibrils (PFFs) accumulated significantly fewer α-syn aggregates than wild-type littermates in the substantia nigra, cortex and hippocampus. HS mouse mutants also displayed lower levels of phosphorylated α-syn in the brain, decreased dopaminergic neuronal loss, and fewer motor deficits than wild-type controls. Moreover, molecular imaging of radiolabeled α-syn PFFs with positron emission tomography (PET)/ computed tomography (CT) revealed faster transport through the brain and spinal cord in mice with less sulfated neuronal HS. Together, these findings provide functional evidence of neuronal HS as a key host cofactor that accelerates α-syn pathology by promoting neuronal internalization and slowing the interstitial fluid (ISF) transport and glymphatic clearance of α-syn fibrils.

## Results

### α-Syn fibrils bind heparin electrostatically

HS colocalizes with Aβ and prion deposits in Alzheimer’s disease and Creutzfeldt-Jakob disease patients and accelerates the kinetics of these disorders in respective mouse models^22–26,34^. To assess whether HSPGs are associated with α-syn inclusions in human disease, we performed dual immunofluorescence for HS and phosphorylated α-syn (pS129-α-syn) aggregates in the temporal cortex of DLB patients and age-matched healthy controls. We observed prominent HS immunoreactivity within the LB of DLB patients, but no intracellular accumulation of HS in healthy controls (Fig. 1a). Moreover, we found HS in the dopaminergic α-syn inclusions of mice 19 weeks after the striatal injections with mouse α-syn PFFs, but not in mice injected with α-syn monomers (Fig. 1b). These findings indicate that HS is enriched within pathological α-syn aggregates in human and mouse brain.

**Figure 1.**
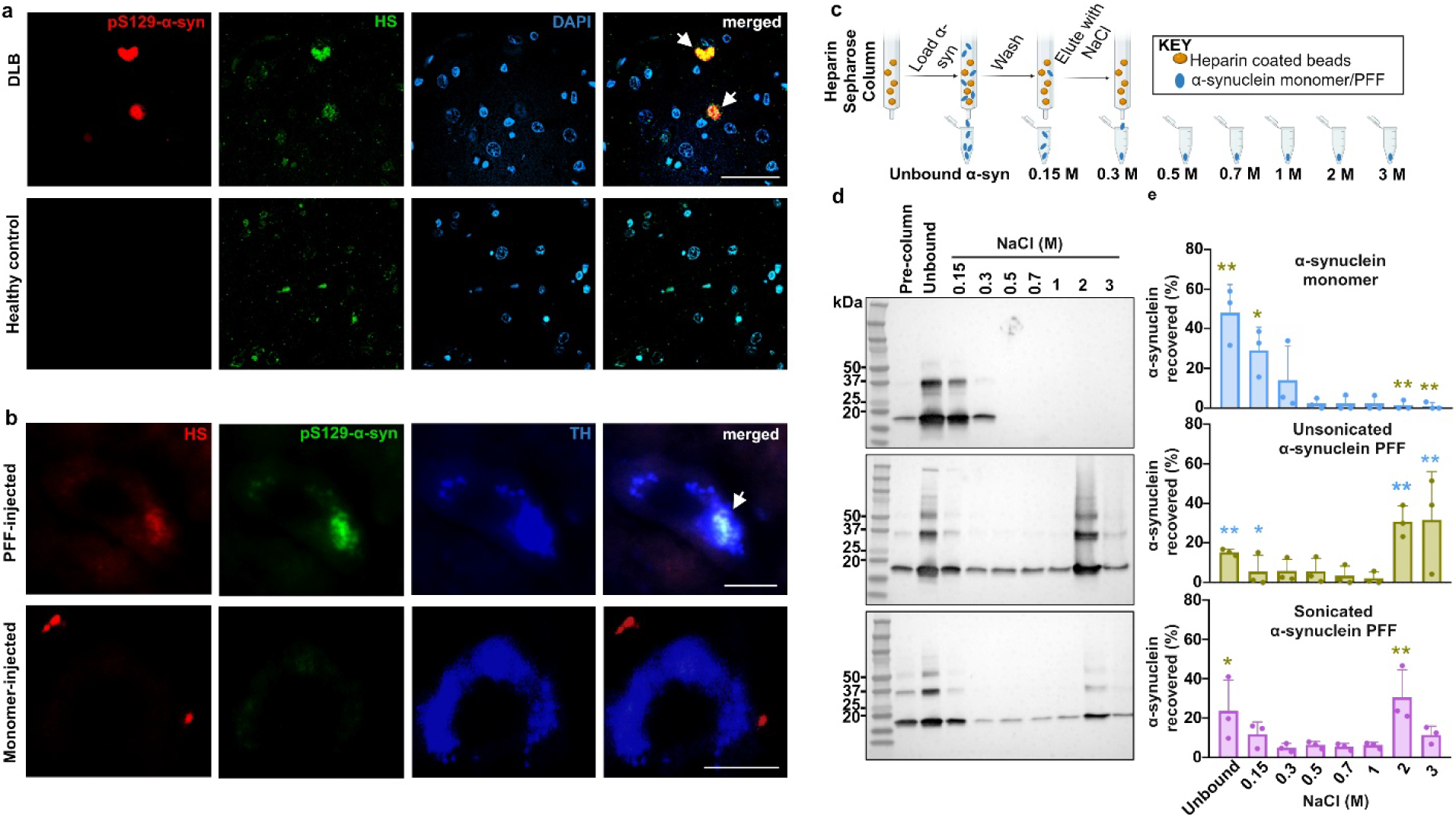
HSPGs bind α-synuclein fibrils with higher affinity than monomers. **(a)** Dual immunofluorescence for HS and phosphorylated α-syn (pS129-α-syn) aggregates in the temporal cortical sections from dementia with Lewy bodies (LB) patients (N = 3). Images show HS associated with pS129-α-syn in LB (white arrows), but was absent in age-matched healthy control tissue. Scale bar = 50 μm. **(b)** Representative images of HS in the dopaminergic LB (white arrows) of PFF-injected mice (N = 3). Scale bar = 10 μm. **(c)** Schematic of heparin-Sepharose affinity chromatography workflow modified from^23^. Recombinant α-syn monomer, unsonicated PFFs (N = 3 independent experiments) or sonicated, short PFFs (N = 3 independent experiments) were applied onto heparin-coated beads, and bound protein was eluted across a stepwise NaCl buffer gradient (150 mM - 3 M). Immunoblots using an antibody to total α-syn were performed. **(d, e)** Western blot (d) and quantifications (e) of the total α-syn in the elution fractions from heparin-Sepharose chromatography. Bar graphs with individual data points show mean ± SD. \**P* < 0.05, \*\**P* < 0.01. Two-way ANOVA with Bonferroni posthoc test (monomer vs PFF; sonicated PFF vs unsonicated PFF).

To assess the capacity of fibrillar α-syn to bind HSPGs, we next tested the binding affinity of heparin, a highly sulfated mimetic of HS polysaccharide chains, to mouse α-syn PFFs. Unsonicated recombinant α-syn monomers and PFFs (average radius of > 100 nm) were applied separately to heparin-Sepharose columns, and heparin-bound protein was eluted across increasing concentrations of NaCl buffers ranging from 0.15 M to 3 M (Fig. 1c). Monomeric α-syn showed little retention on the column (48 ± 14 % of protein recovered in the flow through), with an elution peak at 0.15 M NaCl (29 ± 12 %), indicating weak electrostatic interaction with heparin (Fig. 1d, e). In contrast, PFFs show significantly higher binding affinity for heparin, with only 15 ± 1.3 % protein recovered in the flow through, and a prominent elution peak observed at 3 M NaCl (PFF: 32 ± 25 % and monomer: 1.0 ± 1.8 %). These results suggest that α-syn fibrils, but not monomers, bind to HS with high affinity.

Long PFFs might experience aggregation on the heparin-Sepharose column or have non-specific binding sites that make them difficult to elute. α-Syn PFFs are routinely sonicated to produce shorter fragments that can be more readily internalized by cells while retaining their ability to act as seeds for further α-syn aggregation^35^. Hence, we next applied sonicated (with a 20-30 nm average radius) and unsonicated fibrils (with > 100 nm average radius) onto the heparin-Sepharose columns. Sonicated PFFs displayed strong and not significantly different binding for heparin compared to unsonicated PFFs, with the majority of the protein eluting at 2M NaCl for sonicated PFFs (31 ± 14 %) compared to 3M NaCl for unsonicated PFFs (32 ± 25 %) (Fig. 1d, e). Collectively, our in vitro studies show that α-syn PFFs have unusually high binding capacity for heparin and suggest that the sonicating process routinely performed before the injection in mice or incubation of cells with PFFs would not majorly affect the ability of the α-syn fibrils to bind HS in cultured cells and mouse brain.

### Sulfated HSPGs mediate the neuronal internalization of α-syn fibrils

HSPGs promote the internalization of α-syn fibrils in mouse hippocampal neurons^31^ and iPSC-derived dopaminergic neurons^32^. To test the requirement of HS chains for α-syn fibril internalization in cortical neurons, which show Lewy pathology in PD and DLB patients^36–38^, neurons were isolated from the cortex of wild-type P0 pups, treated with a cocktail of heparinases I, II, and III (5 mU/ml) to enzymatically remove cell surface HS, and incubated with ATTO 488-conjugated α-syn PFFs for 30 minutes (min). The ATTO 488 fluorescence signal of extracellular, not incorporated PFFs was quenched with trypan blue and normalized over the area covered by neurons. Heparinase treatment resulted in a significant reduction in PFF uptake compared with untreated neurons (percentage area covered by PFF signal in untreated neurons: 0.28 ± 0.05 % vs heparinase-treated neurons: 0.03 ± 0.01 %), while no significant difference was observed at 0 min (untreated: 0.011 ± 0.003 % vs heparinase-treated: 0.01 ± 0.003) as shown by confocal imaging and quantitative analysis of the 488-fluorescence signal (Fig. 2a, b). □ ^39^^35^□), significantly reduced fibril uptake compared with untreated controls at 30 min (untreated: 0.26 ± 0.07 % vs chlorate-treated: 0.02 ± 0.005 %), while no significant difference was observed at 0 min (untreated: 0.02 ± 0.005 % vs chlorate-treated: 0.02 ± 0.01 %) (Fig. 2c, d), which suggests that HS sulfation is necessary for the neuronal uptake of α-syn fibrils.

**Figure 2.**
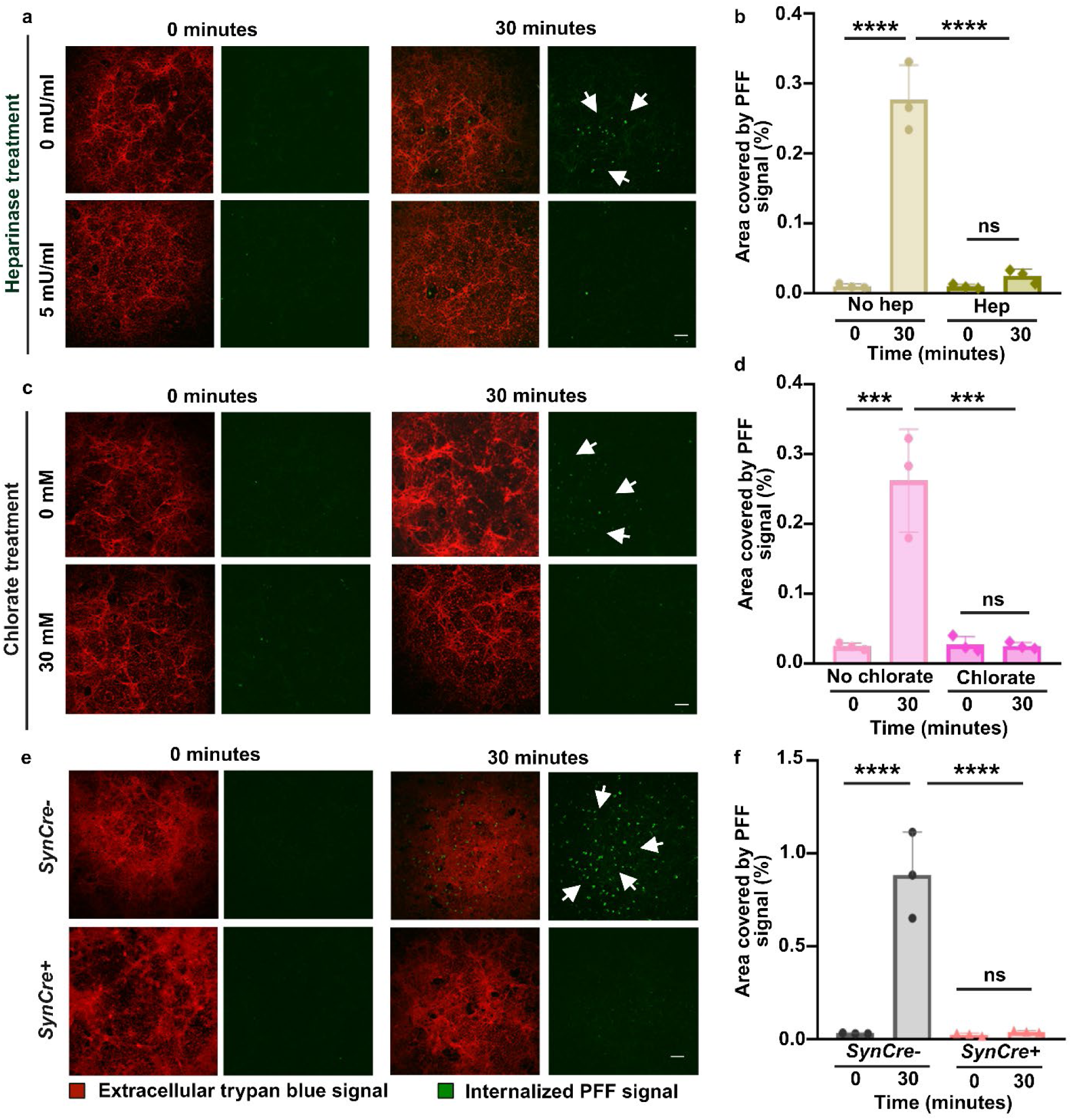
Reducing HSPG sulfation impairs neuronal incorporation of α-syn fibrils. **(a)** Representative confocal fluorescence images of primary cortical neurons from wild-type mice treated with heparinases to remove cell-surface HSPGs prior to incubation with 488-tagged α-syn PFFs. **(b)** Quantification of intracellular 488 fluorescence shows a significant reduction of PFF incorporation in heparinase-treated neurons compared to untreated neurons (N = 3 independent experiments). **(c, d)** Chlorate treatment significantly reduces fibril uptake compared with untreated neurons. **(e, f)** PFF uptake is significantly impaired in the neurons from mice that express lower neuronal HS sulfation (*Ndst1SynCre+*) when compared to wild-type neurons (*Ndst1SynCre-*). In all the incorporation studies, the ATTO 488 fluorescence signal of extracellular, not incorporated PFFs was quenched with trypan blue which resulted in red signal. Scale bar = 100 μm. *SynCre-*and *SynCre+* denote *Ndst1SynCre-* and *Ndst1SynCre+* mice, respectively. Bar graphs with individual data points show mean ± SD. One-way ANOVA with multiple-comparisons and Tukey’s posthoc test. \*\*\**P*<0.001, \*\*\*\**P*<0.0001. Note: trypan blue quenches the fluorescence of extracellular PFFs, even those bound to cellular receptors, thus delineating the neuronal surfaces. Internalized PFFs (green) are indicated by white arrowheads in panels: a, c, e.

To further define the requirement for HS sulfation in neuronal fibril uptake, we next assessed α-syn incorporation in cultured neurons from the *Ndst1SynCre+* mice that express 33 % less sulfation per disaccharide and reduced N-, 2-O- and 6-O-sulfation in the brain than wild-type mice due to conditional knockout of the *Ndst1* gene in neurons with the *Synapsin* 1 (*Syn*) promoter^23^ (Sup. Fig. 1). Again, fibril uptake was almost abrogated in sulfation-deficient neurons at 30 min (*Ndst1SynCre-*: 0.88 ± 0.23 % vs *Ndst1SynCre+* neurons: 0.04 ± 0.01 %), while no significant difference was observed at 0 min (*Ndst1SynCre*-: 0.03 ± 0.00 % vs *Ndst1SynCre+* neurons: 0.03 ± 0.01 %) (Fig. 2 e, f). Taken together, the enzymatic (heparinase), chemical (chlorate), and genetic (*Ndst1SynCre*) approaches provide evidence that sulfated HSPGs are required for efficient neuronal incorporation of α-syn fibrils in vitro.

### Reducing neuronal HS sulfation decreases α-syn pathology in mice

Since HSPGs are major receptors for the neuronal internalization of α-syn fibrils and seem to bind α-syn through the sulfate groups in their chains, we next investigated the role of HSPGs in α-syn pathology in vivo using *Ndst1SynCre* mice and the PFF-induced synucleinopathy model^35,40^. The expression of the *Ndst1* gene in the dopaminergic and the glutamatergic neurons - two neuronal populations particularly vulnerable to α-syn pathology^36–38^ - was first validated with fluorescent in situ hybridization of wild-type mouse brain tissue (Sup. Fig. 2a). Further characterization of the *Ndst1SynCre* mouse model showed that, despite expressing 33 % lower HS sulfation in mouse brain^23^, *Ndst1SynCre+* mice have similar levels of α-syn expression in the brain to those of wild-type littermates (*Ndst1SynCre-*mice) (Sup. Fig. 2b, c).

18 weeks after intrastriatal injections of mouse PFFs, immunofluorescence staining for pS129-α-syn revealed fibril-induced pathology in the brains of *Ndst1SynCre+* and *NdstSynCre-*mice and no pathology in monomer-injected mice (Fig. 3a-f). Notably, *Ndst1SynCre+* mice showed ∼49 % decreased fibril-induced aggregate accumulation in the dopaminergic neurons of the substantia nigra pars compacta (SNpc) (area covered by pS129-α-syn in *Ndst1SynCre-*: 8.5 ± 3.4 vs *Ndst1SynCre+*: 4.4 ± 2.5), ∼60% in the cortex (*Ndst1SynCre-*: 62.7 ± 34.0 vs *Ndst1SynCre+*: 25.4 ± 19.7), and ∼77% in the hippocampus (*Ndst1SynCre-*: 9.2 ± 6.5 vs *Ndst1SynCre+*: 2.1 ± 1.7) relative to *Ndst1SynCre-*littermates (Fig. 3a-f). Serial extraction of α-syn into Triton X-100-soluble and insoluble fractions confirmed an approximately 59% decrease in aggregated α-syn burden in PFF-injected *Ndst1SynCre+* brains compared with *Ndst1SynCre-*controls (*Ndst1SynCre-:* 1.3 ± 0.4 vs *Ndst1SynCre+:* 0.5 ± 0.5), whereas no α-syn aggregates were detected in the monomer-injected brains (*Ndst1SynCre-:* 0.1 ± 0.1 *vs Ndst1SynCre+*: 0.1 ± 0.1) (Fig. 3g,h). Moreover, the 49% decrease in the deposition of α-syn aggregates in SNpc corresponded with a ∼59% increase in dopaminergic neuronal survival as measured with stereology (neuronal count by stereology, *Ndst1SynCre-*: 3,845 ± 824 vs *Ndst1SynCre+*:6,112 ± 1,295) (Fig. 3i, j). None of the brains of monomer-injected *Ndst1SynCre-* and *Ndst1SynCre+* mice showed α-syn aggregation or neuronal loss, and neither genotype showed behavioral deficits (Sup. Fig. 3). Altogether, these transmission studies indicate that neuronal HS is a major driver of fibril-induced α-syn aggregate accumulation and associated dopaminergic neuronal loss in vivo.

**Figure 3.**
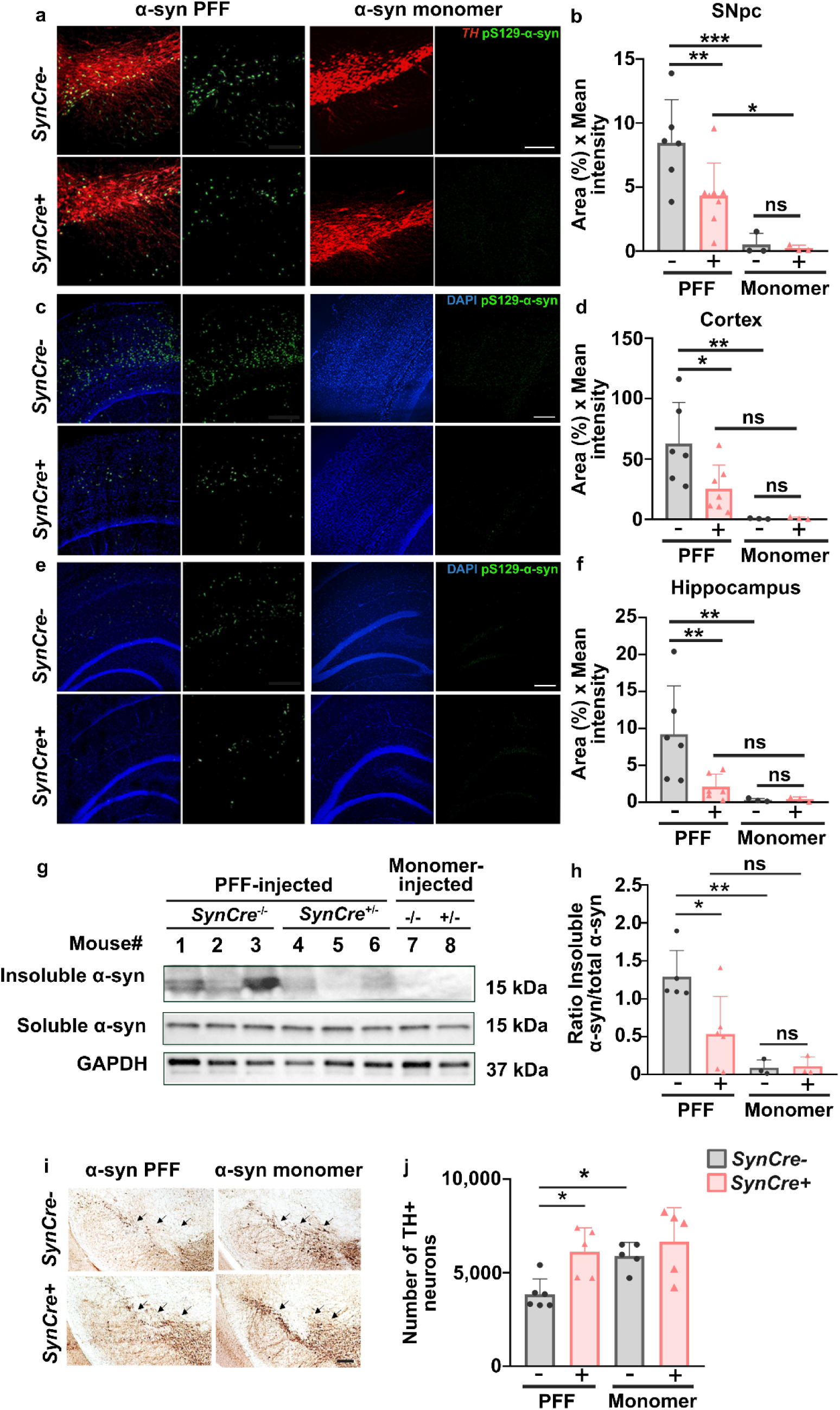
Reduced neuronal HS sulfation decreases fibril-induced α-synuclein accumulation and dopaminergic neuronal loss in vivo. (a, **b)** Representative immunofluorescence images and quantification of pS129-α-syn pathology in the substantia nigra pars compacta (SNpc) of *Ndst1SynCre-* and *Ndst1SynCre+* mice 18 weeks after intrastriatal injection of α-syn PFFs or monomers. *Ndst1SynCre+* mice show an approximately 49 % reduction in pS129-α-syn-positive area compared with *Ndst1SynCre-*littermates. (c, d) Representative images and quantification of pS129-α-syn pathology in the cortex reveal an approximately 60 % reduction in *Ndst1SynCre+* mice. (e, f) Representative images and quantification of pS129-α-syn pathology in the hippocampus show an approximately 77 % reduction in *Ndst1SynCre+* mice. No pS129-α-syn pathology was detected in monomer-injected mice. (g, h) Serial extraction of α-syn into Triton X-100-soluble and proteinase K-treated Triton X-100-insoluble fractions demonstrate a lower 59 % burden of aggregated α-syn in *Ndst1SynCre+* brains compared with *Ndst1SynCre-*controls. (i, j) Representative tyrosine hydrolase (TH) immunostaining and stereological quantification of dopaminergic neurons in the SNpc. *Ndst1SynCre+* mice show a 59 % increase in dopaminergic neuronal survival compared with *Ndst1SynCre-*littermates. *SynCre-* and *SynCre+* denote *Ndst1SynCre-* and *Ndst1SynCre+* mice, respectively. Scale bars = 100 μm. Bar graphs with individual data points show mean ± SD. One-way ANOVA with multiple-comparisons and uncorrected fisher’s LSD posthoc test. \**P*<0.05, **P<0.01, \*\*\**P*<0.001, \*\*\*\**P*< 0.0001.

### Reducing neuronal HS sulfation slows the development of locomotion impairment in PFF-injected mice

To determine whether differences in α-syn pathology were associated with behavioral alterations, PFF- and monomer-injected *Ndst1SynCre+* and *Ndst1SynCre-*mice underwent a battery of behavioral assays assessing locomotion, anxiety-like behavior, fear learning and memory. Open field test, elevated plus maze test, and contextual fear conditioning revealed no differences in the locomotor activity, motor coordination, anxiety-like behavior, and associative memory between genotypes at 12 weeks of age (Sup. Fig 2d-o), suggesting that reducing neuronal HS sulfation does not significantly impact α-syn expression PFF or behavior of synucleinopathies in mice at this young age. However, at 17 weeks post-injection, PFF-injected *Ndst1SynCre+* mice showed increased locomotor activity compared to *Ndst1SynCre-*littermates in the open field test, with significantly longer distance traveled (Fig. 4b) (*Ndst1SynCre-*: 2,529 ± 463 cm vs *Ndst1SynCre+*: 3,800 ± 838 cm), and higher average velocity (Fig. 4c) (*Ndst1SynCre-:* 4.2 ± 0.8 cm/s vs *Ndst1SynCre+*: 6.4 ± 1.4 cm/s) over the 10 min trial. Time spent in the center of the arena did not differ between genotypes (Fig. 4d) (*Ndst1SynCre-*: 109 ± 77 s vs *Ndst1SynCre+*: 96 ± 54 s), suggesting no change in anxiety-like behavior in this assay. These results indicate that while neuronal *Ndst1* deficiency does not seem to alter anxiety levels, it significantly modulates the motor phenotype in PFF-injected mice.

**Figure 4.**
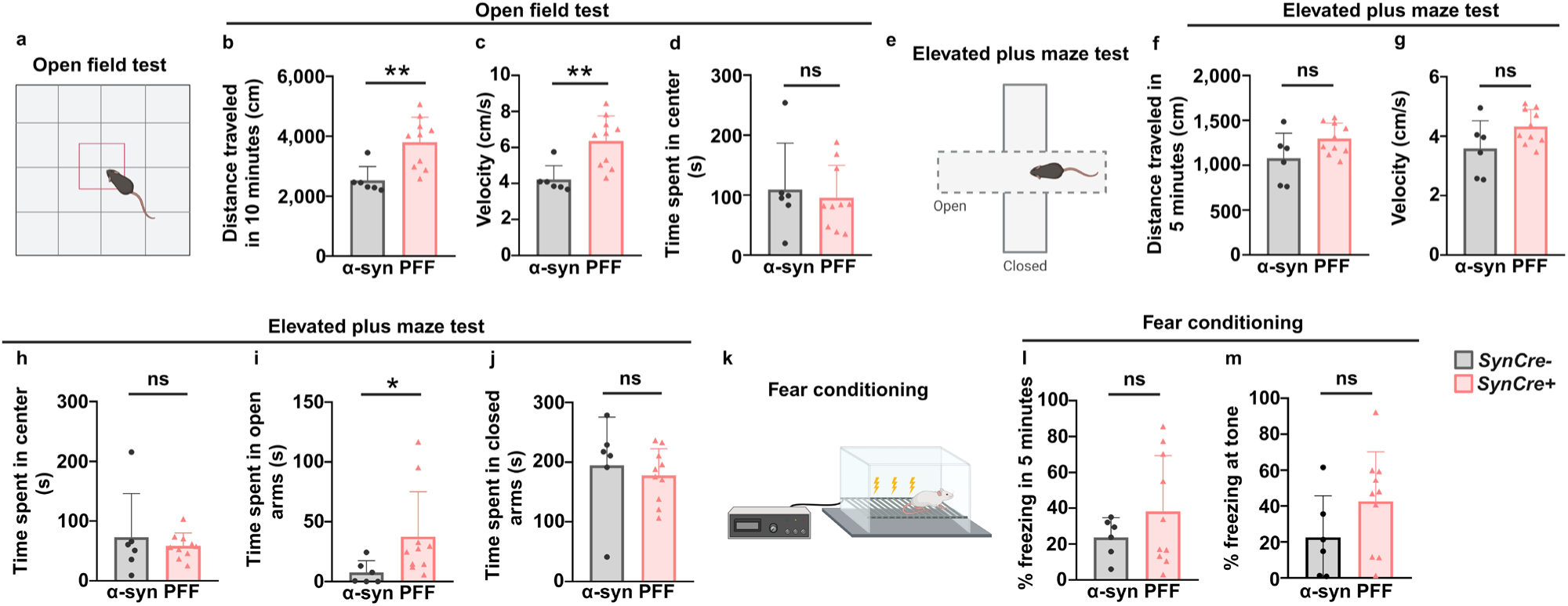
Neuronal HS sulfation deficiency attenuates PFF-induced motor deficits. **(a)** Representative open-field activity schematic testing paradigm performed 17 weeks post PFF injection. **(b, c)** *Ndst1SynCre+* mice show significantly increased total distance traveled and average velocity compared with *Ndst1SynCre-*littermates during the 10-minute trial. **(d)** Time spent in the center of the arena did not differ between genotypes. **(e)** Representative elevated plus-maze schematic testing platform. **(f–h)** Total distance traveled, average velocity, and time spent in the center did not differ between PFF-injected *Ndst1SynCre-* and *Ndst1SynCre+* mice. **(i, j)** *Ndst1SynCre+* mice spent more time in the open arms and similar time in the closed arms than Ndst1SynCre-littermates; however, genotype-dependent differences were observed in monomer-injected controls (Sup. Fig. 3 g,h), suggesting these effects may not be specific to PFF-induced pathology. **(k)** Schematic of the fear-conditioning paradigm. **(l, m)** Freezing during contextual and cued fear-memory testing did not differ between PFF-injected *Ndst1SynCre-* and *Ndst1SynCre+* mice. *SynCre-* and *SynCre+* denote *Ndst1SynCre-* and *Ndst1SynCre+* mice respectively. Bar graphs with individual data points show mean ± SD. Unpaired Welch’s t-test. \**P*<0.05, **P<0.01.

In the elevated plus maze test, there were no differences between PFF-injected genotypes in total distance traveled (Fig. 4f) (*Ndst1SynCre-*: 1077 ± 280 cm vs *Ndst1SynCre+*: 1,296 ± 175 cm), velocity (Fig. 4g) (*Ndst1SynCre-*: 3.6 ± 0.93 cm/s vs *Ndst1SynCre+*: 4.3 ± 0.58 cm/s), or time spent in the center (Fig. 4h) (*Ndst1SynCre-*: 73 ± 73 s vs *Ndst1SynCre+*: 58 ± 22 s). Differences in time spent in the open arms (Fig. 4i) (*Ndst1SynCre-:* 7.6 ± 9.8 s vs *Ndst1SynCre+*: 38 ± 38 s) and closed arms (Fig. 4j) (*Ndst1SynCre-*: 195 ± 81 s vs *Ndst1SynCre+:* 178 ± 45 s) were observed, however, a similar effect was also present in monomer-injected controls (Sup. Fig 3g: *Ndst1SynCre-:* 27 ± 20 s vs *Ndst1SynCre+:* 58 ± 30 s and Sup. Fig 3h: *Ndst1SynCre-:* 219 ± 20 s vs *Ndst1SynCre+:* 164 ± 46 s), suggesting that this change is not specific to PFF-induced pathology, and does not reflect a consistent change in anxiety-related behavior.

To assess associative learning and memory, mice underwent fear conditioning to examine hippocampus-dependent contextual memory and amygdala-dependent cued learning. During contextual testing, freezing behavior was similar between *Ndst1SynCre-* vs *Ndst1SynCre+* mice (Fig. 4l) (Freezing time (%): *Ndst1SynCre-*: 24 ± 11 vs *Ndst1SynCre+*: 38 ± 31), despite a significant decrease in hippocampal α-syn pathology in *Ndst1SynCre+* animals, indicating that changes in hippocampal pathology may not accompany corresponding behavioral alterations in contextual fear performance at the 17 weeks post-injection timepoint. Similarly, cued fear conditioning revealed no differences in freezing during tone presentation (Fig.4m) (Freezing time (%): *Ndst1SynCre-:* 23 ± 23 vs *Ndst1SynCre+:* 43 ± 28). Collectively, these behavioral studies show that neuronal deletion of *Ndst1* reduces PFF-induced motor dysfunction, with no significant effects on anxiety-related behaviors or fear memory.

### Neuronal HS is an obstacle for the interstitial diffusion of the α-syn fibrils

Neuronal HS slows the transport through the brain and the CSF clearance of prion monomers, proteins with a medium binding affinity for heparin^22,23^. To test how neuronal HS modulates ISF transport and glymphatic clearance of α-syn fibrillary aggregates, we quantified the distribution of radiolabeled α-syn PFFs in the brain and spine from *Ndst1SynCre*- and *Ndst1SynCre*+ mice over 72 hours (hr) following intrastriatal injections. α-Syn PFFs were conjugated with the bifunctional chelator deferoxamine maleimide, sonicated to 30-40 nm radius, and radiolabeled with zirconium-89 ([^89^Zr]-^89^Zr]PFFs), a positron-emitting radionuclide with a half-life of 78.4 hr. [^89^Zr]-PFFs were injected under stereotaxic conditions into the right dorsal striatum of *Ndst1SynCre*- and *Ndst1SynCre*+ mice (Fig. 5a). Full body [^89^Zr]-PFFs PET imaging was acquired every 24 hr, followed by computed tomography (CT) for anatomical reference. The brain and spine distribution of the radiolabeled fibrils was quantified to extract metrics of standardized uptake values (SUVs). After completion of imaging at 72 hr, mice were sacrificed and the presence of radioactivity across neurological (brain and spine) and non-neurological (blood, liver, kidneys, spleen, stomach, cecum, leg muscle, femur, and tail) tissues was quantified with a gamma counter.

**Figure 5.**
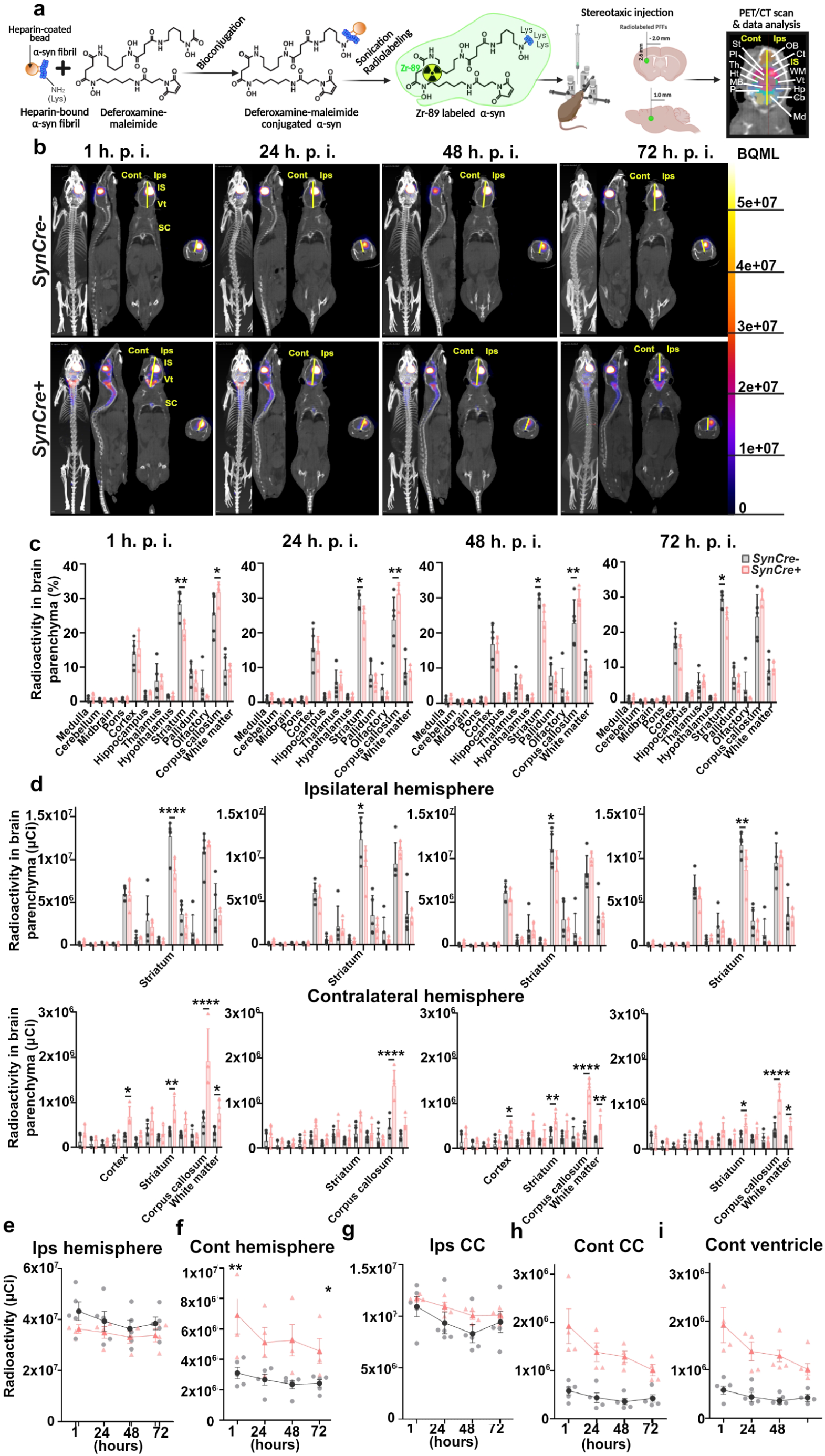
Sulfated HS is an obstacle for the transport of α-synuclein fibrils. **(a)** Schematic of experimental workflow: α-Syn PFFs were conjugated to deferoxamine maleimide (DFO), radiolabeled with zirconium-89 ([^89^Zr]PFFs) and injected under stereotaxic conditions into the right dorsolateral striatum of *Ndst1SynCre-* and *Ndst1SynCre+* mice. PET/CT imaging was performed at 1, 24, 48, and 72 hr after injection. Images were reconstructed and the areas of interest detected with a mouse brain atlas: striatum (St), pallidum (Pl), thalamus (Th), hypothalamus (Ht), midbrain (MB), pons (P), olfactory bulb (OB), cortex (CT), injection site (IS), white matter (WM), ventricle (Vt), hippocampus (Hp), cerebellum (Cb) and medulla (Md). **(b)** Representative PET/CT images showing the distribution of [^89^Zr]PFF in the brain and spinal cord of both genotypes at indicated time points demonstrate distribution through time between *Ndst1SynCre-* and *Ndst1SynCre+* mice (contralateral and ipsilateral hemispheres, cont and ips, respectively). **(c)** Regional distribution of radioactivity in brain parenchyma 1 hr post-injection, expressed as a percentage of total radioactivity in the whole brain regions in both genotypes. **(d)** Quantification of radioactivity (μCi) in the ipsilateral striatum and in the contralateral striatum, cortex, corpus callosum, and white matter 1 hr after injection in both genotypes. **(e, f)** Radioactivity (μCi) retained in the ipsilateral (e) and contralateral (f) hemispheres over time in both genotypes. **(g, h)** Quantification of radioactivity (μCi) retained in ipsilateral corpus callosum (g) and contralateral corpus callosum (h) of both genotypes over time. **(i)** Quantification of radioactivity (μCi) retained in contralateral ventricles of both genotypes over time. Bar graphs with individual data points show mean ± SD. Longitudinal PET/CT data were analyzed using a mixed-effects model, including the genotype × time interaction. Ex vivo measurements were analyzed using One-way ANOVA with multiple comparisons. \**P*<0.05, \*\**P*<0.01, \*\*\*\**P*< 0.0001.

One hr post-injection with [^89^Zr]-PFFs in the dorsolateral striatum, the PET/CT scan showed heightened uptake largely concentrated at the injection site (striatum) and the ipsilateral corpus callosum of *Ndst1SynCre*- and *Ndst1SynCre*+ mice compared to the contralateral brain tissue (Fig. 5b-c, percentage radioactivity in striatum over radioactivity in brain parenchyma). Quantification across brain regions was analyzed using a brain atlas. Notably, the percentage of radioactivity in the striatum of *Ndst1SynCre*+ mice was significantly lower than in the *Ndst1SynCre*-mice (Fig. 5b-c, *Ndst1SynCre*-: 28 ± 4.0 % vs *Ndst1SynCre*+: 21 ± 3.0 %), while the percentage of radioactivity in the corpus callosum of *Ndst1SynCre*+ mice was significantly higher than that of *Ndst1SynCre*-mice (Fig. 5b-c, *Ndst1SynCre*-: 25 ± 6.0 % vs *Ndst1SynCre*+: 32 ± 3.0 %), suggesting that reducing neuronal HS accelerates the transport of [^89^Zr]PFFs from the injection site to neighboring brain areas.

To further understand how neuronal HS controls the kinetics of PFF spread through the central nervous system, we compared the distribution of the radioactivity in the injected (ipsilateral) and the contralateral hemispheres. One hr post-injection with [^89^Zr]PFFs, *Ndst1SynCre-*mice had most of the radioactivity concentrated in the injection site as expected (Fig. 5d, ipsilateral striatum: 30 ± 3.0 % of all the radioactivity in the parenchyma of the right brain hemisphere), ipsilateral corpus callosum (26 ± 5.0 %) and cortex (14 ± 3.0 %), with less than 10% of the radioactivity in the contralateral brain parenchyma (Sup Fig. 4b, c, 5.4 ± 2.0 %). In the *Ndst1SynCre+* ipsilateral hemispheres, however, most of the radioactivity concentrated in the corpus callosum (33 ± 2.0 %), followed by the striatum (23 ± 2.0 %) and cortex (16 ± 4.0 %) and 11 ± 2.5 % of radioactivity spread to the contralateral hemisphere (Fig. 5d and Sup. Fig. 4b, c). *Ndst1SynCre*+ mice showed not only significantly lower radioactive signal at the injection point at the 1 hr post-injection timepoint (Fig. 5d, *Ndst1SynCre*-: 8.3×10^6^ ± 1.7×10^6^ µCi vs *Ndst1SynCre*+: 1.3×10^7^ ± 2.0×10^6^ µCi), but more than two fold higher levels of [^89^Zr]PFFs in the contralateral hemisphere (Fig. 5f, *Ndst1SynCre*-: 3.1×10^6^ ± 6.9×10^5^ µCi vs *Ndst1SynCre*+: 6.9×10^6^ ± 2.0×10^6^ µCi) and more than threefold higher levels in the contralateral corpus callosum (Fig. 5g, *Ndst1SynCre*-: 5.8×10^5^ ± 1.9×10^5^ µCi *vs Ndst1SynCre*+: 1.9×10^6^ ± 7.2×10^5^ µCi), further suggesting that neuronal HS modulates the interstitial transport of α-syn PFFs.

The percentage of radioactivity retention at the injection site did not significantly vary between groups after the first hr post-PFF injection in either *Ndst1SynCre*- or *Ndst1SynCre*+ mice (Sup. Fig. 4d, percent of radioactive signal in the ipsilateral striatum over the radioactivity in the neurological tissues, fixed genotype effect). However, the levels of radioactivity in the ipsilateral hemisphere and corpus callosum slowly but significantly decreased over time for both genotypes (Fig. 5e, g, fixed time effect). Strikingly, the radioactivity in the contralateral hemisphere and corpus callosum decreased significantly faster in the *Ndst1SynCre*+ mice than in the *Ndst1SynCre*-mice (Fig. 5f, time and a genotype fixed effects and mixed time x genotype effect), potentially in line with the higher CSF clearance in the *Ndst1SynCre*+ contralateral ventricles (Fig. 5h, *Ndst1SynCre*-: 21 ± 10 % vs *Ndst1SynCre*+: 27 ± 4.0 %) and the spine+ventricles (Sup. Fig. 4e). Collectively, these PET/CT studies suggest that neuronal HS regulates the interstitial transport of α-syn fibrils, which may in turn impact their clearance to and from the CSF.

### Reducing neuronal HS accelerates the interstitial transport and glymphatic clearance of the α-syn fibrils

The volume and rate of intrastriatal infusion define the initial, mechanically-driven distribution of tracer within the parenchyma^41,42^, and studies of solute transport in mouse striatum typically infuse ∼1µL of volume at a rate of 0.1 µL/min ^43,44^. Even under those conditions, the volume of tracer delivered and dispersed varies several-fold between animals and physiological states, such that comparisons require adjustment for initially delivered dose/volume^44^. Our [^89^Zr]PFF infusions (3 µL at 0.67 µL/min) exceeded these standard parameters, impeding an accurate assessment of the ISF transport of molecules in physiological conditions. We therefore used the 1 hr post-injection image as an animal-specific measure of the infusion-determined distribution, and subtracted it from the 24, 28, and 72 hr images, so that the reported signal reflects redistribution occurring after the acute infusion phase. These calculations confirmed the faster transport of radioactivity through the contralateral CC of *Ndst1SynCre*+ mice (Fig. 6a and b, fixed and mixed genotype and time effects). Moreover, a radical change in the clearance of [^89^Zr]PFFs was observed 24 hr post-injection. While the PFF clearance decreased over time in *Ndst1SynCre*-mice, it increased in the *Ndst1SynCre*+ mice, confirming that reducing neuronal HS sulfation accelerates the ISF transport and CSF clearance of PFFs (Fig. 6b, mixed genotype and time effect).

**Figure 6.**
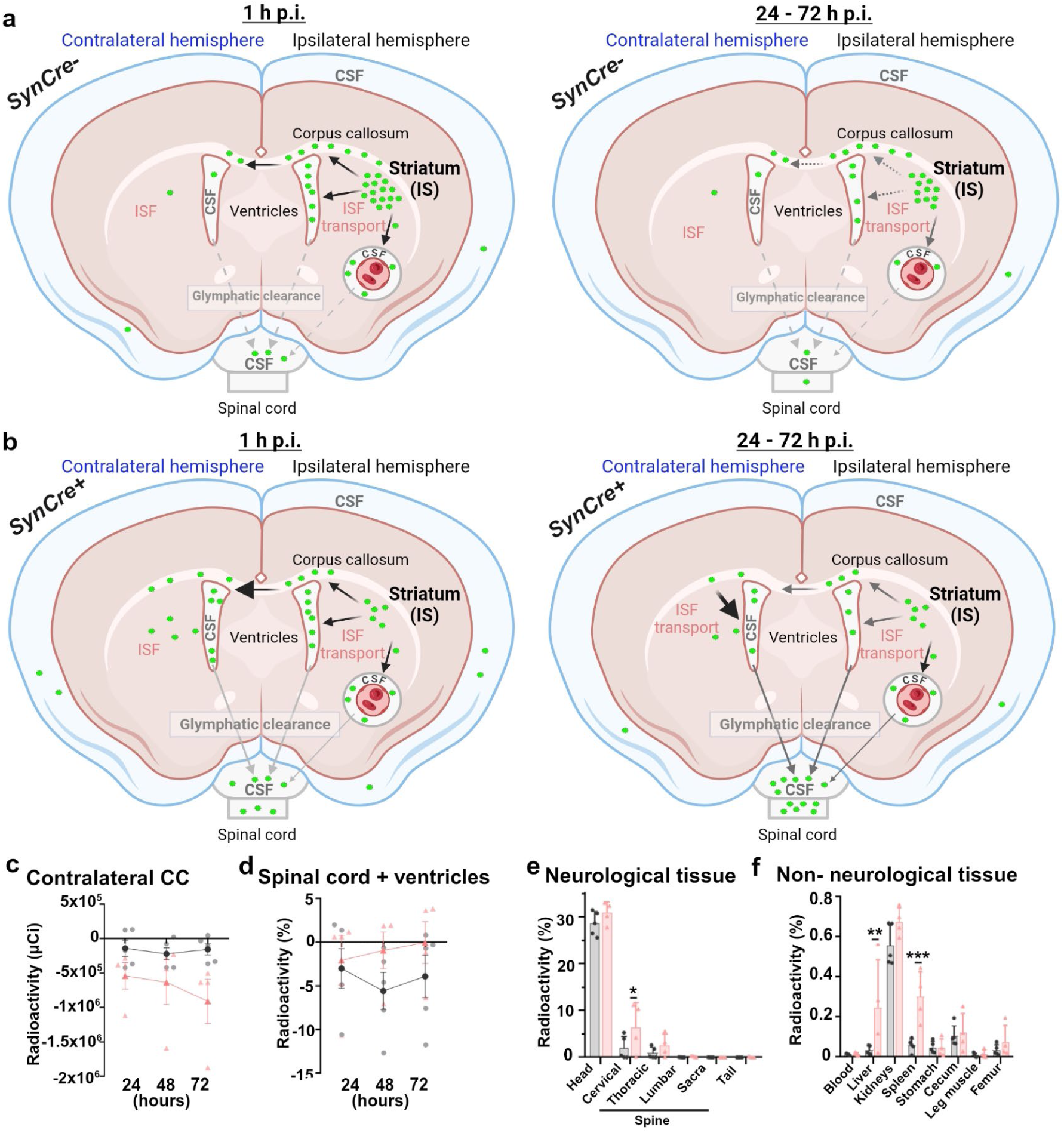
Reduced neuronal HS sulfation accelerates the interstitial transport and clearance of α-synuclein fibrils. (a, **b)** Schematic of the distribution of [^89^Zr]PFFs (green circles) in *Ndst1SynCre-*(a) and *Ndst1Syncre+* (b) mice after their stereotaxic injection in the dorsal striatum (injection site, IS). **(b)** Representative PET/CT images showing the distribution of [^89^Zr]PFF in the brain and spinal cord of both genotypes at indicated time points demonstrate distribution through time between *Ndst1SynCre-* and *Ndst1SynCre+* mice. **(c, d)** Changes in [^89^Zr]PFF-associated radioactivity at 24, 48, and 72 hr after subtraction of the corresponding 1-hr signal, showing transport through the contralateral corpus callosum (c) and spinal cord+ventricles over time (d). **(e,f)** Ex vivo biodistribution of radioactivity at 72 hr of neurological tissue (e) and non-neurological tissue (f), measured by gamma counting and expressed as the percentage of injected dose per gram of tissue (%ID/g). *SynCre-* and *SynCre+* denote *Ndst1SynCre-* and *Ndst1SynCre+* mice, respectively. Bar graphs with individual data points show mean ± SD. Longitudinal PET/CT data were analyzed using a mixed-effects model, including the genotype × time interaction. Ex vivo measurements were analyzed using One-way ANOVA with multiple comparisons. \**P*<0.05, \*\**P*<0.01, \*\*\**P*<0.001, \*\*\*\**P*< 0.0001.

CSF is ultimately cleared to the blood and thus, we assessed how neuronal HS sulfation impacts the spread of PFFs in non-neurological tissues (blood, liver, kidneys, spleen, stomach, cecum, leg muscle, femur, and tail). The biodistribution after necropsy revealed the highest presence of radioactivity (measured as the percent of the injected dose per gram of tissue, %ID/g) in the brain and skull, followed by the upper spine (cervical and thoracic spine sections) of *Ndst1SynCre+* and *Ndst1SynCre-*mice (Fig. 6c). Interestingly, *Ndst1SynCre+* mice showed significantly higher levels of radioactivity in the cervical spine (*Ndst1SynCre*-: 2.0 ± 3.0 % vs *Ndst1SynCre+*: 6.0 ± 5.0 %), liver, and spleen compared to that in *Ndst1SynCre-*mice, (Fig. 6c, d) which further suggests that decreasing neuronal HS sulfation accelerates the clearance of α-syn fibrils. Altogether, these imaging results suggest that neuronal HS might also contribute to α-syn pathology by slowing the ISF transport and CSF clearance of α-syn fibrils.

## DISCUSSION

Mounting studies indicate that HS is a key receptor for the internalization of α-syn fibrils in cultured cells, yet the role of HS in the disease progression of α-syn pathology in vivo is poorly known. Here we establish that the neuronal-derived HS glycosaminoglycans are major modulators of the synucleinopathy progression. We show that α-syn fibrils interact with the sulfate groups in the HS chains with high affinity, and that genetically reducing the sulfation of neuronal HS drastically reduces the accumulation of α-syn aggregates in multiple brain areas, the loss of dopaminergic neurons, and the development of a behavioral phenotype in PFF-injected mice. We also found that reducing HS sulfation accelerates the transport of α-syn fibrils towards CSF clearance and blocks the internalization of α-syn PFFs in cortical neurons. Collectively, our results support a model in which neuronal HS accelerates α-syn pathology by promoting the neuronal internalization and slowing the glymphatic clearance of α-syn aggregates.

HS is present in the extracellular amyloid plaques of Alzheimer’s disease, cerebral amyloid angiopathy, and Creutzfeldt-Jakob disease patients^22,24,25,45^, and its roles as a cellular receptor and cofactor for the fibril formation of α-syn, tau, amyloid-beta (Aβ), and prion fibrils^20,34,46–56^ are well known. This study shows for the first time HS in the α-syn intraneuronal deposits and provides in vivo evidence that neuronal HS acts as a host cofactor that also promotes α-syn pathology. Mice expressing less sulfated neuronal HS showed up to 77% fewer α-syn inclusions in multiple brain areas, and almost 60% less pS129-α-syn in the brain than wild-type mice 18 weeks after injection with PFFs. Importantly, reducing neuronal HS sulfation also protected against dopaminergic neuronal loss and the development of motor impairment, in line with previous findings in PFF-fed *Caenorhabditis elegans* with shorter or less sulfated HS^57^. Our results resemble the reduction in the plaque deposition of Aβ and prions in mouse models of Alzheimer’s disease and some prion disorders with less sulfated neuronal HS^23,26^. Whether modifying HS composition also decreases neurotoxicity and slows the development of cognitive and motor impairment in these disorders is unknown. However, Alzheimer’s disease and prion-infected mice showed longer survival times and/ or less neuroinflammatory response, providing more aspects where HS could contribute to neurodegeneration.

The high binding affinity of α-syn PFFs to heparin and the almost complete blockage of the neuronal incorporation of α-syn PFFs in neurons genetically, enzymatically and chemically modified to express lower HS levels or lower HS sulfation, suggests that neuronal HS would potentially accelerate α-syn pathology by favoring the neuronal internalization of α-syn fibrils. This molecular mechanism might not seem relevant for proteinopathies where the aggregated protein solely accumulates intracellularly and is mainly transported cell-to-cell along axons and neuronal synapses, as in tauopathies and synucleinopathies. Nevertheless, α-syn and tau fibrils are leaked to the extracellular space from dead neurons or actively secreted through vesicular and non-vesicular pathways^58,59^, and are found in CSF and/or blood of dementia with Lewy bodies, Parkinson’s disease, and Alzheimer’s disease patients^8–12,60,61^. Therefore, tackling the cell-to-cell spread of α-syn fibrils by impairing their neuronal internalization appears as a rational therapeutic approach for synucleinopathies. In fact, multiple receptors such as LAG3 or LRP1 promote the uptake of α-syn PFFs in cultured cells and neuronal toxicity in mice^62,63^. However, the expression of some of these receptors in distinct neuronal populations, their subcellular location, and their interactions with α-syn fibrils, and the off-target effects and compensatory signaling are not well known^64,65^, precluding their use as therapeutic targets. HS is an attractive therapeutic target for synucleinopathies for multiple reasons. HSPGs are not only located on the cell surface of neurons, but also in the presynapses^19^, where they might interact with α-syn fibrils and contribute to their trans-synaptic spread. Besides, HS’s binding properties to a desired ligand can be modulated by multiple strategies (for example, heparin mimetics, HS-binding peptides, competitive glycan analogs, enzymes that modify HS structure) with minimal impact on its vast interactome and ubiquitous biology. Importantly, our studies show that *Ndst1SynCre* + mice did not display major modifications in their physiological behavior or in the levels of α-syn expressions, which informs about a novel molecular target for developing therapies against synucleinopathies: the Ndst1 enzyme.

Neuronal HS, and in particular the Ndst1 enzyme, could also be targeted to decrease the α-syn burden in the brain of PD and DLB patients by enhancing their glymphatic clearance. Neuronal HSPGs are cleaved from the plasma membrane by proteases or secreted into the extracellular space in vesicles, where they get inserted into the hyaluronic acid matrix^66^. The negatively charged nature of the HS chains makes them molecular obstacles for the ISF diffusion of some molecules, which potentially triggers the scaffolding of parenchymal and vascular amyloid plaques in Alzheimer’s disease, cerebral amyloid angiopathy, and Creutzfeldt-Jakob disease patients^22,23^. We previously found that decreasing neuronal HS sulfation accelerates the glymphatic clearance of monomeric prions^23^. In this study, the live imaging experiments with radiolabeled α-syn PFFs suggest that neuronal HS also modulates the transport of the α-syn PFFs through the brain extracellular space, which in turn impacts glymphatic clearance to the CSF and then clearance from the CSF compartment itself. This conclusion is supported by i) the significantly higher radioactivity in the hemisphere, corpus callosum and ventricle contralateral to the injection site in the *Ndst1SynCre*+ mice compared with wild-type *Ndst1SynCre*-controls during the first hr post-injection with PFFs; ii) the faster spread through the contralateral corpus callosum and CSF clearance of the [^89^Zr]PFFs in *Ndst1SynCre+* brains during the following 72 hr of the PET scan study; and iii) the higher levels of radioactivity in the cervical spine, spleen and kidneys of *Ndst1SynCre+* mice at necropsy. Collectively, these studies provide a framework for investigating the role of neuronal HS, or other components of the brain extracellular matrix, in the glymphatic clearance of pathogenic α-syn, a mechanism that is impaired in the brains of PD and DLB disease patients^67,68^.

Our [^89^Zr]-PFFs PET imaging studies also yielded the first detailed description of the in vivo distribution of the α-syn PFFs after injection in the dorsolateral striatum, a commonly used technique to investigate mechanisms of α-syn pathology. The strikingly low percentage of radioactivity retained in the injection site of *Ndst1SynCre-* and *Ndst1SynCre+* mice during the first hr post PFF injection (approximately 22 and 14 %, respectively), together with the high percentage that was detected in their ventricles and spinal cord at this early time point (approximately 25 %), suggests that the fast infusion rates of the PFFs solution (0.67 µl/min) might have accelerated the ISF bulk flow in the mouse brain (estimated at 0.04-0.13 µl/min in the brain of healthy adult rat^53^) towards CSF clearance. The most remarkable differences in the ISF transport and CSF clearance between the two genotypes also occurred during this first hr, which might be a result of the lower resistance to the PFF infusion provided by the brain parenchyma with less sulfated HS. The striatal injections of serial dilutions of PFFs in wild-type mice showed that a 10-fold and a 40-fold dilution are needed to reach the 70 % and 50 % loss in α-syn accumulation registered in the cortex and the SN of *Ndst1SynCre+* mice^54^. Thus, it is unlikely that the 30 % lower retention of PFFs at the injection site (estimated from the PET scan imaging at 1 hr post-injection with [^89^Zr]PFFs) is responsible for the delay in α-syn deposition in *Ndst1SynCre+* mice. Nevertheless, these PET/CT studies prompt the infusion of the PFFs at a lower rate to prevent the artefactual exacerbation of certain phenotypes.

In summary, our study provides evidence that neuronal HS controls the progression of α-syn pathology potentially by promoting neuronal internalization and slowing glymphatic clearance of pathogenic α-syn. The HS-mediated internalization of α-syn fibrils in dopaminergic neurons from PD patients^28^, and the detection of HS in the α-syn deposits of DLB patients shown in this study, support the development of therapeutics to prevent the interaction between neuronal HS and α-syn fibrils to slow the progression of synucleinopathies. Further studies will show whether the decreased cell-to-cell spread, α-syn deposition, and dopaminergic neuronal loss impact the development of motor, cognitive, and psychiatric abnormalities in later disease stages.

## MATERIALS AND METHODS

### Ethics statement

All animal procedures were approved by the Institutional Animal Care and Use Committee (IACUC) at the University of Alabama at Birmingham (protocol number: 22908) and were conducted in accordance with the National Institutes of Health Guide for the Care and Use of Laboratory Animals. Mice were housed under standard conditions on a 12 hr light/dark cycle with *ad libitum* access to food and water and were routinely monitored for health and welfare concerns.

*Ndst1^f/f^* mice^69^ were crossed with *Synapsin*-*Cre* transgenic mice to drive neuron-specific recombination. Offspring carrying the *Synapsin*-*Cre* transgene and two floxed alleles were used as *Ndst1^f/f^SynCre^+/−^*(*SynCre+)* and *Ndst1^f/f^SynCre^−/−^* (*SynCre-)* mice, respectively. Both male and female mice were used for experiments.

### Preparation of recombinant α-syn preformed fibrils

α-Syn preformed fibrils (PFFs) were generated from recombinant monomeric α-syn following a previously published protocol^70^. Briefly, α-syn monomers were expressed in E. coli, purified using ion exchange and size exclusion chromatography, and verified for purity by SDS-PAGE before fibril assembly. Endotoxin was removed using a Pierce endotoxin removal spin column, and endotoxin in monomeric α-syn was 0.076 EU/µg. α-Syn monomers were incubated with 150 mM KCl and 50 mM Tris-HCl at 37°C and agitated for 7 days at 1,000 rpm. Fibrils were later isolated from the monomers by centrifugation for 10 min at 13,200 rpm, their protein concentration was determined by spectrophotometer, and a buffer consisting of 50 mM Tris-HCl and 150 mM KCl was used to bring the concentration to 5 mg/ml. PFFs were stored at −80°C until use. Immediately before usage, PFFs were sonicated for 15 min (amplitude 37; pulse on 3 s; pulse off 2 s) using a Qsonica sonicator with circulating water at 15°C. The fragmentation and radius of PFFs was confirmed using dynamic light scattering (Wyatt Technology).

### Fluorescent *in situ* hybridization

Fluorescent *in situ* hybridization (FISH) was performed on 20 µm fixed coronal brain sections from wild-type mice using the RNAscope Multiplex Fluorescent v2 Assay (Advanced Cell Diagnostics, Cat No. 323100) according to the manufacturer’s instructions. Probes targeting *TH*, *Ndst1*, and *Glut1* (*Slc2a1*) were hybridized and visualized using multiplex fluorophores (Opal 520, Alexa Fluor 555 / Opal 570, and Opal 690 / Cy5). Nuclei were counterstained with DAPI, and slides were mounted using ProLong Gold Antifade Mountant. Images were captured on a Nikon C2 confocal microscope using an air objective lens. Standard channel settings were used to acquire fluorescence signals for DAPI and probe-specific fluorophores. Image files were exported and processed using NIS-Elements software.

### Intrastriatal injection of recombinant α-syn fibrils and tissue sampling

Three-months-old mice (N=6-11/group) were anesthetized with vaporized isoflurane and secured on a stereotaxic frame (Kopf Instruments). Animals received bilateral stereotaxic injections of sonicated α-syn preformed fibrils (PFFs) or monomeric α-syn into the dorsolateral striatum (+1.0 mm AP and ±2.0 mm ML from bregma;-2.6 mm DV from dura). Each hemisphere was injected with 2 µL of α-syn (5 mg/mL; 10 µg total protein per mouse) at a constant infusion rate of 0.5 µL/min. Following completion of the injection, the needle was left in place for 4 min before being gradually withdrawn to minimize reflux. Mice were administered with 10 units of 1.5 mg/ml carprofen subcutaneously immediately before and the following day after surgery.

At 18 weeks post-injection, PFF-injected and α-syn monomer injected control mice were made unconscious with CO_2_ and perfused transcardially with 1x phosphate-buffered saline (PBS). During necropsy, brains were dissected into halves: one hemisphere was fixed in 4% paraformaldehyde (PFA) and cryoprotected in 30% sucrose in PBS, while the contralateral hemisphere was snap-frozen for immunoblotting analysis. Fixed hemispheres were sectioned coronally at 40 µm thickness using a Leica SM2010 R freezing microtome. Serial sections were collected in six-well plates, ensuring each well contained slices spaced 240 μm apart, representing the entire mouse brain. Sections were stored at −20 °C in cryoprotectant solution (50% glycerol, 0.01% sodium azide in Tris-buffered saline) until use.

### Brain samples from dementia with Lewy bodies patients

The postmortem brain tissue sample from a patient with DLB and a healthy control were used to assess the presence of HS in pS129-α-syn aggregates. The brain samples were provided by the Emory University Goizueta Alzheimer’s Disease Research Center (GADRC). A more detailed description of the demographic information and pathological scoring can be found in ^71^ (samples DLB-1 and Control-1).

### Immunofluorescence and immunohistochemistry

For immunofluorescence, free-floating brain sections were washed with cold 1x Tris-buffered saline (TBS), followed by antigen retrieval in 10 mM sodium citrate buffer (pH 6.0) at 37 °C for 1 hr. Sections were rewashed in TBS and blocked for 1 hr at 4°C in 10% normal goat serum (NGS) containing 0.1% Triton X-100 in 1× TBS. Primary antibodies: rabbit anti-phosphorylated α-syn (Abcam, ab51253; 1:500) and chicken anti-tyrosine hydroxylase (Abcam, ab76442; 1:500) were diluted in 10% NGS in 1× TBS and incubated with sections for 24 hr at 4 °C. Following washes in 1× TBS, sections were incubated with secondary antibodies: Cy3-conjugated goat anti-rabbit IgG (Jackson ImmunoResearch, 111-165-144; 1:500) and Cy5-conjugated goat anti-chicken IgY (Thermo Fisher Scientific, A21449; 1:500) for 2 hr at 4 °C in 10% NGS in 1× TBS. After secondary incubation, sections were washed and counterstained with DAPI (Sigma aldrich, 1:4,000 in 1× TBS) for 5 min at room temperature (RT), followed by final washes in TBS. Sections were then mounted onto charged glass slides, air-dried, and coverslipped using ProLong Gold antifade mounting medium (Invitrogen). All incubations and wash steps were performed with gentle agitation on an orbital shaker. Human DLB brain tissue sections were treated with TrueBlack Lipofuscin Autofluorescence Quencher (Cell Signaling Technology, cat. no. 92401) to reduce lipofuscin autofluorescence following the manufacture’s protocol.

For immunohistochemistry, free-floating brain sections were washed in cold 1× TBS and incubated in 3% hydrogen peroxide for 20 min at RT to quench endogenous peroxidase activity. Sections were rinsed in distilled water followed by antigen retrieval in sodium citrate buffer (pH 9.0) at 37 °C for 30 min. Following washes in TBS, sections were blocked for 1 h at 4 °C in 10% NGS containing 0.3% Triton X-100 in 1× TBS. Sections were incubated with rabbit anti-tyrosine hydroxylase (TH) antibody (Abcam, ab152; 1:2,000) diluted in 1 % NGS with 0.3 % Triton X-100 in TBS for 24 h at 4 °C. After washings in TBS, sections were incubated with peroxidase-conjugated goat anti-rabbit IgG (H+L) secondary antibody (Jackson ImmunoResearch, 115-035-003; 1:500) diluted in 2.5% NGS with 0.3% Triton X-100 in TBS for 4 h at RT. Immunoreactivity was visualized using ImmPACT DAB substrate (Vector Laboratories, SK-4105), developed for approximately 2 min, and the reaction was stopped by rinsing in distilled water. Sections were washed, mounted onto glass slides in 1× TBS, air-dried, and coverslipped using Permount mounting medium (Fisher Scientific, SP15-100).

### Quantification of phosphorylated α-synuclein pathology

Image acquisition and quantification were performed by an investigator blinded to experimental conditions. Brain sections were imaged at 20× magnification using a VS200 slide scanner (Olympus). Images were visualized in QuPath, and quantitative analyses were performed using Fiji/ImageJ.

For each mouse, multiple sections from the cortex, hippocampus, and substantia nigra pars compacta (SNpc) were analyzed. Regions of interest (ROIs) were manually defined in ImageJ using anatomical landmarks. Images were thresholded using fixed, region-specific thresholds applied uniformly across all samples. Phosphorylated α-syn-positive area was quantified within each ROI, and measurements were averaged across sections and hemispheres to generate a single value per mouse for statistical analysis.

### Unbiased stereology

Unbiased stereological quantification of tyrosine hydroxylase (TH+) neurons in the SNpc was performed using an Olympus BX51 brightfield microscope with Stereo Investigator software (MBF Bioscience) and the optical fractionator probe. For each animal, 4–6 coronal sections spanning the SNpc were analyzed using systematic random sampling. Low magnification (4×) was used to delineate SNpc boundaries, and high magnification (40×) was used for neuron counting. Counting was performed using a 50 × 50 µm counting frame, an optical dissector height of 24 µm, and an average section thickness of 33 µm, with guard zones applied. The Gundersen coefficient of error (m = 1) was calculated for each animal and maintained below 0.12.

### Behavior testing

All behavioral experiments were conducted in the UAB FBS Behavioral Core Facility. Mice were transported to the behavior core one week prior to the start of testing to allow acclimatization to the testing environment. On each testing day, animals were allowed to acclimate to the testing room for at least 30 min before the start of the experiment. Mice were given 1–3 days of rest between behavioral tests to minimize stress. All behavioral testing and analysis were performed by two investigators blinded to genotype and treatment. Behavioral apparatuses were cleaned thoroughly between trials using 70% ethanol or 2% chlorhexidine to eliminate olfactory cues.

#### Open field test

General locomotor activity and anxiety-like behavior were assessed using the open field test. Mice were placed individually into a square open field arena (43 × 43 cm) with 30-cm-high opaque walls to prevent visual access to the external environment. Each mouse was allowed to freely explore the arena for 10 min. Locomotor activity was recorded and analyzed using Ethovision XT software (Noldus). Parameters measured included total distance traveled, velocity, and percentage of time spent in the center zone. The center and peripheral zones were defined a priori within the software, and mice were tracked throughout the session.

#### Elevated maze test

Anxiety-like behavior was assessed using the elevated plus maze. The apparatus consisted of two open arms and two closed arms arranged in a plus configuration and elevated 50 cm above the floor. Each arm measured 67 cm, with closed arms enclosed by 15 cm high opaque walls. Mice were placed individually in the center of the maze and allowed to freely explore the apparatus for 5 min. Animal movement was recorded and analyzed using Ethovision XT software (Noldus). Parameters quantified included total distance traveled, velocity, and time spent in open arms, closed arms and center.

#### Fear conditioning

Fear conditioning was performed using VideoFreeze software with a grid floor connected to a shock generator. On the training day, mice were placed individually in the chamber and allowed to explore for 2 min, after which a 30 s auditory tone was played followed by a 2 s mild foot shock (0.2 mA). After a 60 s interval, the tone and the shock pairing were repeated once, and mice remained in the chamber for an additional 60 s before being returned to their home cage. Percent time freezing was calculated as the time spent freezing divided by the total time analyzed. On the day following training, contextual fear memory was assessed by returning mice to the same chamber and recording freezing behavior for 5 min. On day 3, cued fear conditioning was assessed in a modified context by replacing the grid floor with solid inserts and altering visual and olfactory cues. The same auditory tone used during training was presented twice for 30 s, separated by 60 s intervals, without foot shock, and the average percent freezing during tone presentations was used for analysis.

### Primary neuronal cultures

Primary cortical neurons were prepared from P0 *Ndst1SynCre* and WT mice as previously described^72^. Tail biopsies were collected at the time of dissection to confirm genotypes. Cortices were dissected in ice-cold dissection medium supplemented with MgCl₂, CaCl₂, HEPES, and glucose, and meninges were removed. Tissue was enzymatically dissociated with trypsin (Gibco) for 15 min at 37 °C, followed by addition of DNase I (Roche) and gentle triturating. Cell suspensions were filtered through a 70-µm cell strainer and centrifuged to remove debris. Neurons were resuspended in high-glucose culture medium and plated onto poly-L-lysine-coated culture dishes or coverslips. Cultures were maintained at 37 °C with 5% CO₂, fed the day after plating with a half media change, and supplemented with fresh medium on day 3.

### α-Syn internalization assay

Neuronal uptake of PFFs was assessed using an internalization assay with Alexa Fluor 488–labeled α-syn PFFs, as previously described^73^. Primary cortical neurons were incubated with sonicated α-syn-Alexa 488 PFFs (1 µg/mL) in cold PBS at 4 °C for 30 min to allow surface binding, then transferred to 37 °C for 30 min to allow internalization. Extracellular fibrils were quenched with 1 mM trypan blue. For enzymatic disruption of cell-surface heparan sulfate, neurons were pretreated with heparinases I, II, and III (5 mU each) for 30 min prior to fibril exposure. In separate experiments, sulfation was inhibited by treating neurons with 30 mM sodium chlorate for 24 h before addition of fibrils. Images were acquired using a confocal microscope with a 10× objective (excitation/emission: 470/550 nm for fibrils and 560/630 nm for trypan blue). Nine fields per plate were imaged and analyzed using ImageJ. Internalized fibril signal was quantified and normalized to trypan blue fluorescence associated with the neuronal membrane. All experiments were performed in three independent replicates.

### Heparin-Sepharose Chromatography

Heparin-binding properties of α-syn monomers, unsonicated preformed fibrils (PFFs), and sonicated PFFs were assessed by heparin-Sepharose affinity chromatography. α-Syn monomers and unsonicated PFFs were prepared as described above and diluted to the desired concentration in PBS (0.14 mg/ml). Sonicated PFFs were adjusted to 1 mL in PBS and subjected to ultracentrifugation (100,000 × g for 30 min at 4 °C) to separate insoluble fibrillar material from soluble species. The supernatant was collected, and the fibril-enriched pellet was resuspended in PBS; an aliquot was retained as a pre-column. For chromatography, Heparin-Sepharose 6 Fast Flow beads (Cytiva) were packed into Bio-Rad disposable columns and equilibrated with 150 mM NaCl in 25 mM HEPES (pH 7.3). α-Synuclein monomers, unsonicated PFFs, and sonicated PFFs were applied to separate columns, and the flow-through was collected and reapplied two to three times to maximize binding. Columns were washed with equilibration buffer, and bound α-syn species were step-eluted using increasing concentrations of NaCl (150 mM-3 M) in 25 mM HEPES (pH 7.3). Fractions were stored at −80 °C until analysis by immunoblotting. All experiments were performed in three independent replicates.

### Western blot

Mouse brain hemispheres without the olfactory bulbs were manually homogenized and sonicated in lysis buffer (50 mM Tris-HCl pH 7.4, 175 mM NaCl, and 5 mM EDTA pH 8.0) with 1% Triton X-100. Samples were incubated on ice for 30 min before being centrifuged for 60 min at 15,000 × g to extract Triton X-100-soluble proteins suspended in the supernatant. Pellets were then resuspended in lysis buffer containing 2% SDS and sonicated to extract Triton X-100-insoluble proteins. Samples were immunoblotted for α-syn following proteinase K digestion of the SDS-soluble fraction. Protein concentrations were assessed by BCA assay (Thermo Fisher Scientific). Tissue lysates were incubated for 5 min at 95C in 4x sample loading buffer (0.25M Tris-HCl pH 6.8, 8% SDS, 200 mM DTT, 30% glycerol, Bromophenol Blue). Equal amounts of protein (20 μg) were loaded per well, resolved on 4-20% SDS-polyacrylamide gels, and transferred to nitrocellulose membranes. After blocking in 5% nonfat dry milk in TBST (25 mM Tris-HCl, pH 7.6, 137 mM NaCl, 0.05% Tween 20), membranes were incubated overnight at 4°C in primary antibody solution containing Mouse Anti α-syn (BD Transduction Laboratory, 610787; 1:1,000), p-Syn (Abcam, 51253; 1:1000), or Rabbit GAPDH (Cell signaling Technologies, 14C10; 1:1000). Samples were then incubated with HRP-conjugated Goat Anti-mouse or Anti-rabbit secondary antibodies (Jackson ImmunoResearch; 115-035-003; 1:2,000 or Jackson ImmunoResearch; 111-035-003; 1:2,000) for one hr at RT. After washing with TBST, blots were developed with enhanced chemiluminescence (GE Healthcare) and imaged with a ChemiDoc XRS+ (Bio-Rad Laboratories).

### α-Syn PFFs radiolabeling

α-Syn PFFs were buffer-exchanged into 0.1 M carbonate, pH 9, using a 7 kDa MW zeba column. α-Syn PFFs were incubated with p-SCN-Bn-DFO (macrocyclics) for 1 hr at 37°C at a molar ratio of 3:1 DFO to DFO-α-Syn PFFs. After incubation, α-Syn PFFs were buffer-exchanged into 1 M HEPES pH 7 twice to remove residual DFO.

Zirconium-89 [^89^Zr] oxalate was purchased from the UAB Cyclotron Facility. [^89^Zr] was neutralized using 1 M HEPES and 2 M NaOH to a pH of 7 and combined with sonicated DFO-α-Syn PFFs at a specific activity of 30 uCi/ug. Samples were incubated for 1 hr at 37°C and radiochemical purity was evaluated using iTLC paper developed in 50 mM DTPA. Under these conditions, [89Zr]Zr-DFO-α-Syn PFFs remain at the baseline, and free [^89^Zr] moves with the solvent. TLC strips were quantified using an AR2000 scanner (Eckert and Zeigler). Radiochemical purity was >95%, and fibrils were buffer-exchanged into PBS to remove the HEPES and oxalate. The final concentration was 13 µCi/µl.

### PET imaging of [^89^Zr]-PFFs: Stereotaxic injection of [^89^Zr]-PFFs, PET image acquisition and analysis

The stereotaxic injections of mice (15 weeks old) (n = 4-5 mice/genotype) were performed as described above: intrastriatal injection of recombinant α-syn fibrils and tissue sampling section, with some modifications to account for radioactivity. The amount of radioactivity in the Hamilton syringe was measured with a dose calibrator before and after every injection to quantify the amount of [^89^Zr]PFFs injected. Approximately 27 µCi of [^89^Zr]PFFs (Sup. Fig. 4a) were injected in the right hemisphere of the mice (+1.0 mm AP and ±2.0 mm ML from bregma;-2.6 mm DV from dura) at a rate of 670 nl/min. Animals were removed from the stereotaxic device and placed on the preclinical PET/CT scanner (Mediso Nanoscan), and radioactivity measurements were collected at 1 hr, 24, 48, and 72 hrs post-injection. Scans were acquired as a static 20-min PET scan followed by a 5-min computed tomography (CT) scan for anatomical reference. Following the last imaging session, hr mice were sacrificed and a biodistribution of tissue was collected. Radioactivity (%ID/g) was quantified with gamma counter (Perkin Elmer Wizard 2).

PET images were reconstructed (Mediso InterView XP) and processed for quantitative assessment of distribution of whole brain and regional uptake of radioactivity (VivoQuant, Invicro). Regions of interest (ROIs) and brain segmentations were extracted (Brain Atlas, VivoQuant) using CT images.

## Statistical analysis

Statistical analyses were performed using GraphPad Prism 8 software (GraphPad Software, San Diego, CA). Data are presented as mean ± standard deviation (SD). Between-group comparisons were conducted using Student’s t-test for two-group comparisons. For multiple comparisons, one-way analysis of variance (ANOVA) was performed, followed by post-hoc tests to identify specific group differences. Longitudinal PET/CT data were analyzed using a mixed-effects model, including the genotype × time interaction. Statistical significance was defined as a *P*-value < 0.05.

## Acknowledgements

This study was funded by the National Institutes of Health grants R00AG061251 (PA-C), the American Parkinson’s Disease Association (PA-C), the Alzheimer’s of Central Alabama predoctoral scholar fellowship (SD), and project 3 of PPG HL131474 (JDE). The Emory University GADRC is funded by NIH grant P30AG066511. The University of Alabama at Birmingham Preclinical Imaging Shared Facility and Ciclotron Facility are funded by NIH grant P30CA013148.

## Author contributions

Conceptualization: PA-C, SD, LV-D, JI, JDE

Methodology: PA-C, LV-D, SD, GS, PD, RB, MG, CM, SEL, AS, RC

Investigation: SD, PD, RB, JH, RQG, HH, EEH, SLS, MG, GS

Visualization: SD, RQG, HH, EEH, SLS

Funding acquisition: PA-C, SD, JDE, LV-D, MG

Project administration: PA-C

Supervision: PA-C

Writing – original draft: PA-C, SD

Writing – review & editing: All authors.

## References

1 Spillantini, M. G., Crowther, R. A., Jakes, R., Hasegawa, M. & Goedert, M. alpha-Synuclein in filamentous inclusions of Lewy bodies from Parkinson’s disease and dementia with lewy bodies. Proc Natl Acad Sci U S A 95, 6469–6473 (1998). 10.1073/pnas.95.11.6469

2 Volpicelli-Daley, L. A. et al. Exogenous alpha-synuclein fibrils induce Lewy body pathology leading to synaptic dysfunction and neuron death. Neuron 72, 57–71 (2011). 10.1016/j.neuron.2011.08.033

3 Mahul-Mellier, A. L. et al. The process of Lewy body formation, rather than simply alpha-synuclein fibrillization, is one of the major drivers of neurodegeneration. Proc Natl Acad Sci U S A 117, 4971–4982 (2020). 10.1073/pnas.1913904117

4 Angot, E., Steiner, J. A., Hansen, C., Li, J. Y. & Brundin, P. Are synucleinopathies prion-like disorders? Lancet Neurol 9, 1128–1138 (2010). 10.1016/S1474-4422(10)70213-1

5 Luk, K. C. et al. Pathological alpha-synuclein transmission initiates Parkinson-like neurodegeneration in nontransgenic mice. Science 338, 949–953 (2012). 10.1126/science.1227157

6 Uemura, N., Uemura, M. T., Luk, K. C., Lee, V. M. & Trojanowski, J. Q. Cell-to-Cell Transmission of Tau and alpha-Synuclein. Trends Mol Med 26, 936–952 (2020). 10.1016/j.molmed.2020.03.012

7 Schaser, A. J. et al. Trans-synaptic and retrograde axonal spread of Lewy pathology following pre-formed fibril injection in an in vivo A53T alpha-synuclein mouse model of synucleinopathy. Acta Neuropathol Commun 8, 150 (2020). 10.1186/s40478-020-01026-0

8 El-Agnaf, O. M. et al. Alpha-synuclein implicated in Parkinson’s disease is present in extracellular biological fluids, including human plasma. FASEB J 17, 1945–1947 (2003). 10.1096/fj.03-0098fje

9 Lee, H. J., Patel, S. & Lee, S. J. Intravesicular localization and exocytosis of alpha-synuclein and its aggregates. J Neurosci 25, 6016–6024 (2005). 10.1523/JNEUROSCI.0692-05.2005

10 Tokuda, T. et al. Decreased alpha-synuclein in cerebrospinal fluid of aged individuals and subjects with Parkinson’s disease. Biochem Biophys Res Commun 349, 162–166 (2006). 10.1016/j.bbrc.2006.08.024

11 Foulds, P. G. et al. Phosphorylated alpha-synuclein can be detected in blood plasma and is potentially a useful biomarker for Parkinson’s disease. FASEB J 25, 4127–4137 (2011). 10.1096/fj.10-179192

12 Mollenhauer, B. et al. alpha-Synuclein and tau concentrations in cerebrospinal fluid of patients presenting with parkinsonism: a cohort study. Lancet Neurol 10, 230–240 (2011). 10.1016/S1474-4422(11)70014-X

13 Sorrentino, Z. A. et al. Intrastriatal injection of alpha-synuclein can lead to widespread synucleinopathy independent of neuroanatomic connectivity. Mol Neurodegener 12, 40 (2017). 10.1186/s13024-017-0182-z

14 Tran, H. T. et al. Alpha-synuclein immunotherapy blocks uptake and templated propagation of misfolded alpha-synuclein and neurodegeneration. Cell Rep 7, 2054–2065 (2014). 10.1016/j.celrep.2014.05.033

15 Zou, W. et al. Blocking meningeal lymphatic drainage aggravates Parkinson’s disease-like pathology in mice overexpressing mutated alpha-synuclein. Transl Neurodegener 8, 7 (2019). 10.1186/s40035-019-0147-y

16 Cui, H. et al. Decreased AQP4 Expression Aggravates a-Synuclein Pathology in Parkinson’s Disease Mice, Possibly via Impaired Glymphatic Clearance. J Mol Neurosci 71, 2500–2513 (2021). 10.1007/s12031-021-01836-4

17 Fujita, K. et al. Mutant alpha-synuclein propagates via the lymphatic system of the brain in the monomeric state. Cell Rep 42, 112962 (2023). 10.1016/j.celrep.2023.112962

18 Sarrazin, S., Lamanna, W. C. & Esko, J. D. Heparan sulfate proteoglycans. Cold Spring Harb Perspect Biol 3 (2011). 10.1101/cshperspect.a004952

19 Zhang, P. et al. Heparan Sulfate Organizes Neuronal Synapses through Neurexin Partnerships. Cell 174, 1450–1464 e1423 (2018). 10.1016/j.cell.2018.07.002

20 Holmes, B. B. et al. Heparan sulfate proteoglycans mediate internalization and propagation of specific proteopathic seeds. Proc Natl Acad Sci U S A 110, E3138–3147 (2013). 10.1073/pnas.1301440110

21 Stopschinski, B. E. et al. Specific glycosaminoglycan chain length and sulfation patterns are required for cell uptake of tau versus alpha-synuclein and beta-amyloid aggregates. J Biol Chem 293, 10826–10840 (2018). 10.1074/jbc.RA117.000378

22 Aguilar-Calvo, P. et al. Shortening heparan sulfate chains prolongs survival and reduces parenchymal plaques in prion disease caused by mobile, ADAM10-cleaved prions. Acta Neuropathol 139, 527–546 (2020). 10.1007/s00401-019-02085-x

23 Aguilar-Calvo, P. et al. Neuronal Ndst1 depletion accelerates prion protein clearance and slows neurodegeneration in prion infection. PLoS Pathog 19, e1011487 (2023). 10.1371/journal.ppat.1011487

24 Snow, A. D. et al. The presence of heparan sulfate proteoglycans in the neuritic plaques and congophilic angiopathy in Alzheimer’s disease. Am J Pathol 133, 456–463 (1988).

25 Snow, A. D., Kisilevsky, R., Willmer, J., Prusiner, S. B. & DeArmond, S. J. Sulfated glycosaminoglycans in amyloid plaques of prion diseases. Acta Neuropathol 77, 337–342 (1989). 10.1007/BF00687367

26 Liu, C. C. et al. Neuronal heparan sulfates promote amyloid pathology by modulating brain amyloid-beta clearance and aggregation in Alzheimer’s disease. Sci Transl Med 8, 332ra344 (2016). 10.1126/scitranslmed.aad3650

27 Huang, R. et al. Ext1-targeted gene therapy ameliorates tauopathy by inhibiting heparan sulfate-dependent tau propagation. Pharmacol Res 230, 108265 (2026). 10.1016/j.phrs.2026.108265

28 Skaanning, L. K. et al. The Non-Fibrillating N-Terminal of alpha-Synuclein Binds and Co-Fibrillates with Heparin. Biomolecules 10 (2020). 10.3390/biom10081192

29 Mehra, S. et al. Glycosaminoglycans have variable effects on alpha-synuclein aggregation and differentially affect the activities of the resulting amyloid fibrils. J Biol Chem 293, 12975–12991 (2018). 10.1074/jbc.RA118.004267

30 Tao, Y. et al. Heparin induces alpha-synuclein to form new fibril polymorphs with attenuated neuropathology. Nat Commun 13, 4226 (2022). 10.1038/s41467-022-31790-7

31 Zhang, Q. et al. A myosin-7B-dependent endocytosis pathway mediates cellular entry of alpha-synuclein fibrils and polycation-bearing cargos. Proc Natl Acad Sci U S A 117, 10865–10875 (2020). 10.1073/pnas.1918617117

32 Vanderperre, B. et al. Novel regulators of heparan sulfate proteoglycans modulate cellular uptake of alpha-synuclein fibrils. Commun Biol 8, 1426 (2025). 10.1038/s42003-025-08786-2

33 Fiorini, M. R., Li, J., Fon, E. A., Farhan, S. M. K. & Thomas, R. A. Neural networks reveal novel gene signatures in Parkinson disease from single-nuclei transcriptomes. NPJ Parkinsons Dis 11, 304 (2025). 10.1038/s41531-025-01147-0

34 Cotman, S. L., Halfter, W. & Cole, G. J. Agrin binds to beta-amyloid (Abeta), accelerates abeta fibril formation, and is localized to Abeta deposits in Alzheimer’s disease brain. Mol Cell Neurosci 15, 183–198 (2000). 10.1006/mcne.1999.0816

35 Abdelmotilib, H. et al. alpha-Synuclein fibril-induced inclusion spread in rats and mice correlates with dopaminergic Neurodegeneration. Neurobiol Dis 105, 84–98 (2017). 10.1016/j.nbd.2017.05.014

36 Surmeier, D. J., Obeso, J. A. & Halliday, G. M. Selective neuronal vulnerability in Parkinson disease. Nat Rev Neurosci 18, 101–113 (2017). 10.1038/nrn.2016.178

37 Henderson, M. X. et al. Spread of alpha-synuclein pathology through the brain connectome is modulated by selective vulnerability and predicted by network analysis. Nat Neurosci 22, 1248–1257 (2019). 10.1038/s41593-019-0457-5

38 Weber, M. A. et al. Alpha-synuclein pre-formed fibrils injected into prefrontal cortex primarily spread to cortical and subcortical structures and lead to isolated behavioral symptoms. bioRxiv (2023). 10.1101/2023.01.31.526365

39 Keller, K. M., Brauer, P. R. & Keller, J. M. Modulation of cell surface heparan sulfate structure by growth of cells in the presence of chlorate. Biochemistry 28, 8100–8107 (1989). 10.1021/bi00446a021

40 Chung, H. K., Ho, H. A., Perez-Acuna, D. & Lee, S. J. Modeling alpha-Synuclein Propagation with Preformed Fibril Injections. J Mov Disord 13, 77–79 (2020). 10.14802/jmd.19046.e

41 Basser, P. J. Interstitial pressure, volume, and flow during infusion into brain tissue. Microvasc Res 44, 143–165 (1992). 10.1016/0026-2862(92)90077-3

42 Chen, M. Y., Lonser, R. R., Morrison, P. F., Governale, L. S. & Oldfield, E. H. Variables affecting convection-enhanced delivery to the striatum: a systematic examination of rate of infusion, cannula size, infusate concentration, and tissue-cannula sealing time. J Neurosurg 90, 315–320 (1999). 10.3171/jns.1999.90.2.0315

43 Pla, V. et al. A real-time in vivo clearance assay for quantification of glymphatic efflux. Cell Rep 40, 111320 (2022). 10.1016/j.celrep.2022.111320

44 Kroesbergen, E. et al. Glymphatic clearance is enhanced during sleep. bioRxiv (2024). 10.1101/2024.08.24.609514

45 Mayfield, J. E. et al. Multi-omic analysis of meningeal cerebral amyloid angiopathy reveals enrichment of unsubstituted glucosamine and extracellular proteins. J Neuropathol Exp Neurol 84, 398–411 (2025). 10.1093/jnen/nlaf018

46 Goedert, M. et al. Assembly of microtubule-associated protein tau into Alzheimer-like filaments induced by sulphated glycosaminoglycans. Nature 383, 550–553 (1996). 10.1038/383550a0

47 Castillo, G. M., Lukito, W., Wight, T. N. & Snow, A. D. The sulfate moieties of glycosaminoglycans are critical for the enhancement of beta-amyloid protein fibril formation. J Neurochem 72, 1681–1687 (1999). 10.1046/j.1471-4159.1999.721681.x

48 Wong, C. et al. Sulfated glycans and elevated temperature stimulate PrP(Sc)-dependent cell-free formation of protease-resistant prion protein. EMBO J 20, 377–386 (2001). 10.1093/emboj/20.3.377

49 Cohlberg, J. A., Li, J., Uversky, V. N. & Fink, A. L. Heparin and other glycosaminoglycans stimulate the formation of amyloid fibrils from alpha-synuclein in vitro. Biochemistry 41, 1502–1511 (2002). 10.1021/bi011711s

50 Berriman, J. et al. Tau filaments from human brain and from in vitro assembly of recombinant protein show cross-beta structure. Proc Natl Acad Sci U S A 100, 9034–9038 (2003). 10.1073/pnas.1530287100

51 Horonchik, L. et al. Heparan sulfate is a cellular receptor for purified infectious prions. J Biol Chem 280, 17062–17067 (2005). 10.1074/jbc.M500122200

52 Wadia, J. S., Schaller, M., Williamson, R. A. & Dowdy, S. F. Pathologic prion protein infects cells by lipid-raft dependent macropinocytosis. PLoS One 3, e3314 (2008). 10.1371/journal.pone.0003314

53 Bazar, E. & Jelinek, R. Divergent heparin-induced fibrillation pathways of a prion amyloidogenic determinant. Chembiochem 11, 1997–2002 (2010). 10.1002/cbic.201000207

54 Sandwall, E. et al. Heparan sulfate mediates amyloid-beta internalization and cytotoxicity. Glycobiology 20, 533–541 (2010). 10.1093/glycob/cwp205

55 Kanekiyo, T. et al. Heparan sulphate proteoglycan and the low-density lipoprotein receptor-related protein 1 constitute major pathways for neuronal amyloid-beta uptake. J Neurosci 31, 1644–1651 (2011). 10.1523/JNEUROSCI.5491-10.2011

56 Ihse, E. et al. Cellular internalization of alpha-synuclein aggregates by cell surface heparan sulfate depends on aggregate conformation and cell type. Sci Rep 7, 9008 (2017). 10.1038/s41598-017-08720-5

57 Chen, M. et al. Heparan sulfate proteoglycans mediate prion-like alpha-synuclein toxicity in Parkinson’s in vivo models. Life Sci Alliance 5 (2022). 10.26508/lsa.202201366

58 Brahic, M., Bousset, L., Bieri, G., Melki, R. & Gitler, A. D. Axonal transport and secretion of fibrillar forms of alpha-synuclein, Abeta42 peptide and HTTExon 1. Acta Neuropathol 131, 539–548 (2016). 10.1007/s00401-016-1538-0

59 Gustafsson, G. et al. Secretion and Uptake of alpha-Synuclein Via Extracellular Vesicles in Cultured Cells. Cell Mol Neurobiol 38, 1539–1550 (2018). 10.1007/s10571-018-0622-5

60 Emmanouilidou, E. et al. Assessment of alpha-synuclein secretion in mouse and human brain parenchyma. PLoS One 6, e22225 (2011). 10.1371/journal.pone.0022225

61 Saman, S. et al. Exosome-associated tau is secreted in tauopathy models and is selectively phosphorylated in cerebrospinal fluid in early Alzheimer disease. J Biol Chem 287, 3842–3849 (2012). 10.1074/jbc.M111.277061

62 Mao, X. et al. Pathological alpha-synuclein transmission initiated by binding lymphocyte-activation gene 3. Science 353 (2016). 10.1126/science.aah3374

63 Gu, H. et al. Lymphocyte Activation Gene 3 (Lag3) Contributes to alpha-Synucleinopathy in alpha-Synuclein Transgenic Mice. Front Cell Neurosci 15, 656426 (2021). 10.3389/fncel.2021.656426

64 Zhang, Y. et al. An RNA-sequencing transcriptome and splicing database of glia, neurons, and vascular cells of the cerebral cortex. J Neurosci 34, 11929–11947 (2014). 10.1523/JNEUROSCI.1860-14.2014

65 Emmenegger, M. et al. LAG3 is not expressed in human and murine neurons and does not modulate alpha-synucleinopathies. EMBO Mol Med 13, e14745 (2021). 10.15252/emmm.202114745

66 Soles, A. et al. Extracellular Matrix Regulation in Physiology and in Brain Disease. Int J Mol Sci 24 (2023). 10.3390/ijms24087049

67 Ding, X. B. et al. Impaired meningeal lymphatic drainage in patients with idiopathic Parkinson’s disease. Nat Med 27, 411–418 (2021). 10.1038/s41591-020-01198-1

68 Pierobon Mays, G., et al. Reduced cerebrospinal fluid motion in patients with Parkinson’s disease revealed by magnetic resonance imaging with low b-value diffusion weighted imaging. Fluids Barriers CNS 21, 40 (2024). 10.1186/s12987-024-00542-8

69 Grobe, K. et al. Cerebral hypoplasia and craniofacial defects in mice lacking heparan sulfate Ndst1 gene function. Development 132, 3777–3786 (2005). 10.1242/dev.01935

70 Stoyka, L. E. et al. Behavioral defects associated with amygdala and cortical dysfunction in mice with seeded alpha-synuclein inclusions. Neurobiol Dis 134, 104708 (2020). 10.1016/j.nbd.2019.104708

71 Digraskar, S. et al. Formic acid impairs alpha-synuclein seeding activity in human and mouse brains. bioRxiv (2025). 10.1101/2025.07.14.664807

72 Lawrence, J. A. et al. Diminished Neuronal ESCRT-0 Function Exacerbates AMPA Receptor Derangement and Accelerates Prion-Induced Neurodegeneration. J Neurosci 43, 3970–3984 (2023). 10.1523/JNEUROSCI.1878-22.2023

73 Froula, J. M. et al. Defining alpha-synuclein species responsible for Parkinson’s disease phenotypes in mice. J Biol Chem 294, 10392–10406 (2019). 10.1074/jbc.RA119.007743

