## Supplementary data for "Neuronal heparan sulfate controls the brain accumulation and glymphatic clearance of pathogenic α-synuclein"

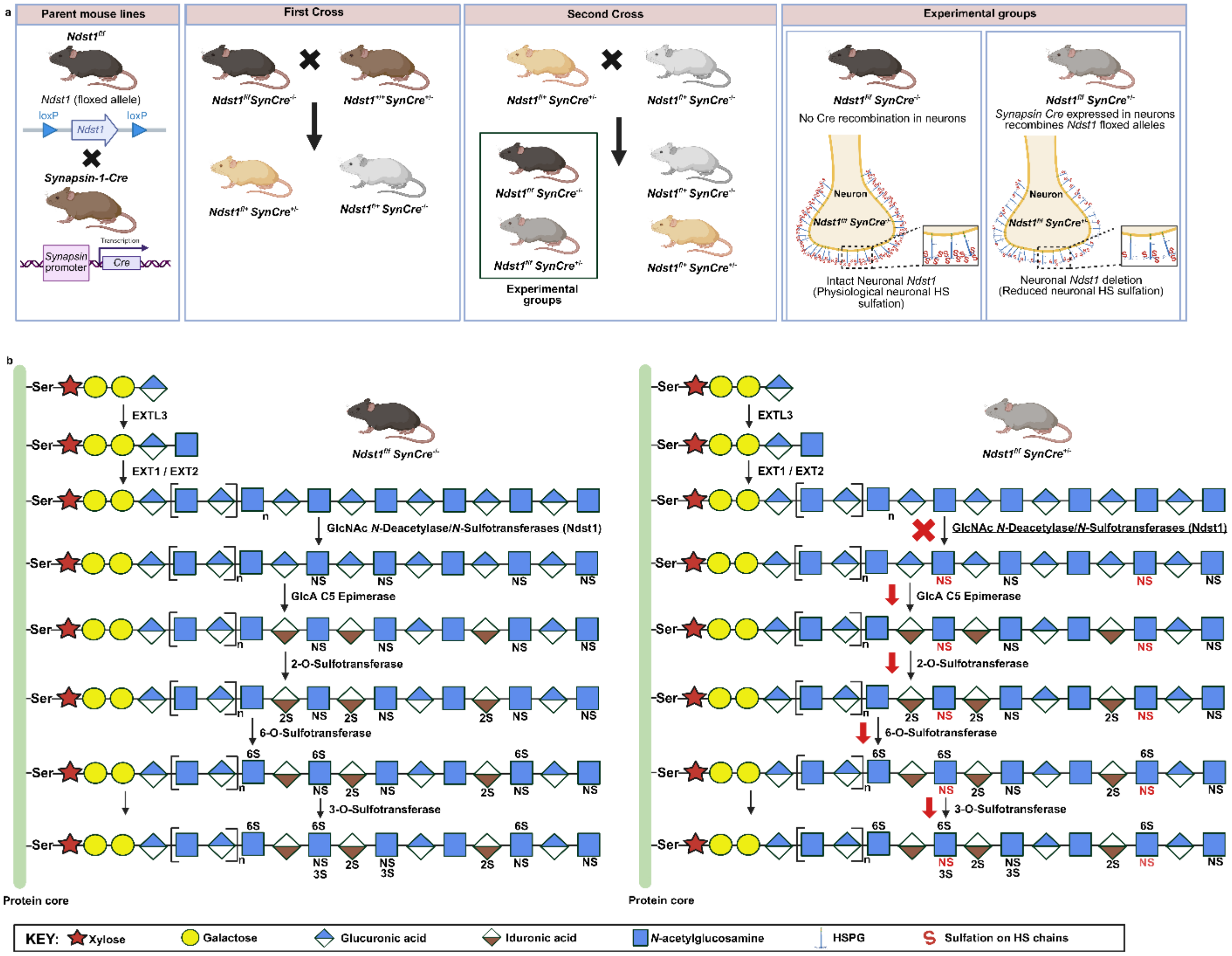


**Sup. Fig. 1. Generation of neuron-specific *Ndst1* conditional knockout mice for targeted disruption of heparan sulfate sulfation. (a)** Breeding strategy for generating neuron-specific *Ndst1* conditional knockout (*Ndst1*$^{f/f}$ SynCre$^{+/-}$) mice and Cre-negative littermate controls (*Ndst1*$^{f/f}$ SynCre$^{-/-}$). Synapsin-1-driven Cre expression mediates neuronal deletion of floxed *Ndst1* alleles decreasing the HS sulfation in these cells. **(b)** Schematic illustration of heparan sulfate (HS) biosynthesis and the predicted effects of neuronal *Ndst1* deletion on HS chain modification. HS chains are synthesized on a serine residue of a core protein through the sequential addition of sugar residues by EXTL3 and the EXT1/EXT2 complex. Ndst1 (*N*-deacetylase/*N*-sulfotransferase 1) catalyzes glucosamine *N*-deacetylation and *N*-sulfation, enabling subsequent modifications by C5-epimerase and 2-*O*-, 6-*O*-, and 3-*O*-sulfotransferases. In

*Ndst1*$^{f/f}$ SynCre$^{-/-}$ control neurons, intact Ndst1 activity supports normal glucosamine *N*-sulfation and subsequent HS modifications. In contrast, neuronal *Ndst1* deletion in *Ndst1*$^{f/f}$ SynCre$^{+/-}$mice is expected to reduce *N*-sulfation and downstream sulfation, as indicated by the red cross and arrows, resulting in HS chains with altered sulfation patterns.

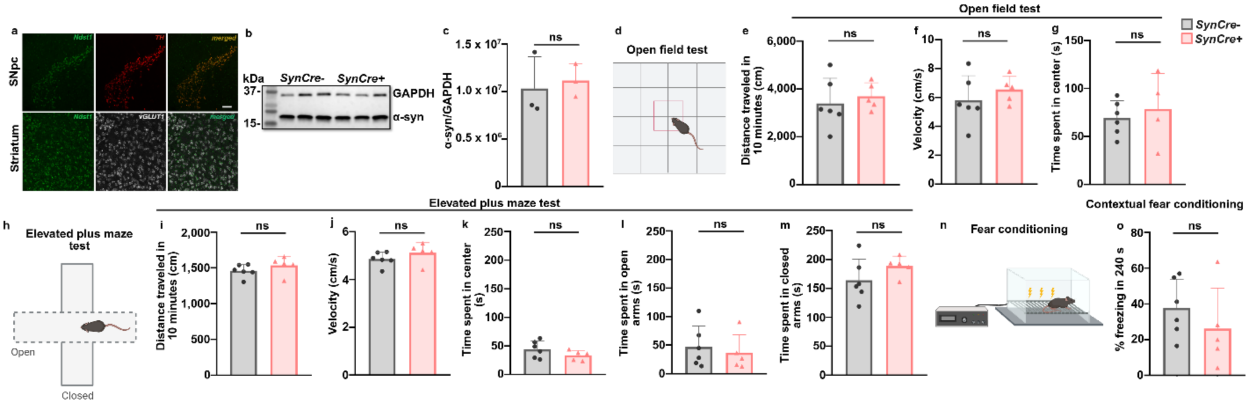


**Sup. Fig. 2. Reduced neuronal HS sulfation does not alter the physiological α-synuclein levels or behavior in mice. (a)** Representative in situ hybridization images showing *Ndst1* gene expression in the substantia nigra pars compacta (SNpc) and striatum. Scale bar = 100 µm. **(b, c)** Representative western blot (b) and densitometric quantification (c) of α-syn levels in brain lysates from 90 days-old *Ndst1SynCre*- and *Ndst1SynCre+* mice. α-Syn levels were normalized to GAPDH and were comparable between genotypes. **(d)** Schematic of the open-field test. (e-g) Distance traveled (cm) (e), velocity (cm/s) (f), and time spent in the center zone (g) during the open-field test. (h) Schematic of the elevated plus-maze test. (i-m) Distance traveled (cm) (i), velocity (cm/s) (j), and time spent in the center (s) (k), open arms (l), and closed arms (m). (n) Schematic of the fear-conditioning paradigm. (o) Contextual fear memory measured as percentage time freezing. No significant differences were observed between Ndst1SynCre- and Ndst1SynCre+ mice. Panels c, e-g, i-m and o: Unpaired t-test.

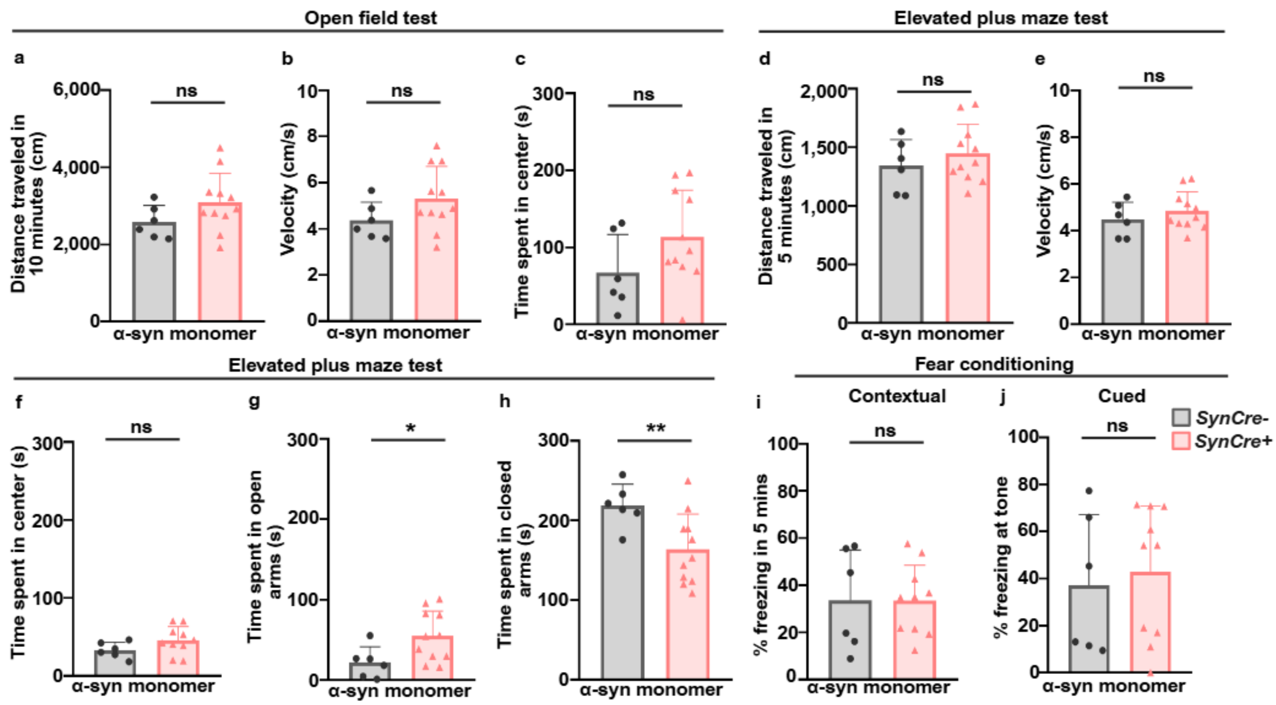


**Sup. Fig. 3. Monomer-injected *Ndst1SynCre-* and *Ndst1SynCre+* mice mostly show comparable behavioral performance. (a-c)** Open-field behavior of monomer-injected mice, including distance traveled (cm) (a), velocity (cm/s) (b), and time spent in the center zone(s) (c) showed no significant differences between genotypes. **(d-h)** Elevated plus-maze behavior, including distance traveled (cm) (d), velocity (cm/s) (e), and time spent in the center (s) (f), open arms (g), and closed arms (h). *Ndst1SynCre-* and *Ndst1SynCre+* mice differed significantly in the time spent in the open and closed arms, whereas the other elevated plus-maze measures were comparable. **(i, j)** Contextual (i) and cued (j) fear memory measured as percentage time freezing. No significant differences were observed between monomer-injected *Ndst1SynCre-* and *Ndst1SynCre+* mice. Unpaired t-test. **P*<0.05, **P<0.01.

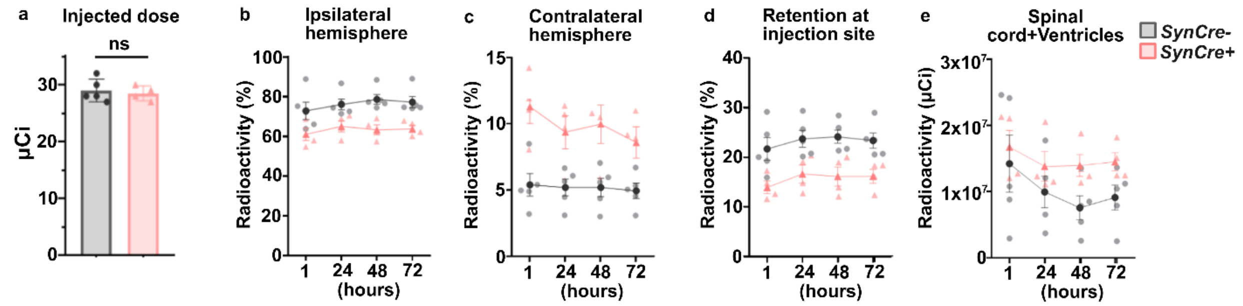


**Sup. Fig. 4. [$^{89}$Zr]PFFs show altered brain and spinal cord distribution in mice with reduced neuronal HS sulfation. (a)** Radioactivity remaining in the syringe after injection, confirming that similar amounts of [$^{89}$Zr]PFFs were administered to *Ndst1SynCre-* and *Ndst1SynCre+* mice. **(b, c)** Distribution of radioactivity in the ipsilateral (c) and contralateral (d) hemispheres over the total radioactive signal in brain and spinal cord across 72 hr. **(d)** Percentage of radioactivity retained at the striatal injection site over the total radioactive signal in brain and spinal cord. **(e)** [$^{89}$Zr]PFFs -associated radioactivity in the spinal cord and cerebral ventricles over 72 hr. Longitudinal data were analyzed using a mixed-effects model with genotype and time as fixed effects and a genotype × time interaction. Data are presented as mean ± SD; points represent individual animals. Panel a: Unpaired t-test.
